# Metabolic potential for reductive acetogenesis and a novel energy-converting [NiFe] hydrogenase in *Bathyarchaeia* from termite guts – a genome-centric analysis

**DOI:** 10.1101/2020.12.10.419648

**Authors:** Hui Qi Loh, Vincent Hervé, Andreas Brune

**Author notes:** **Correspondence:** Andreas Brune.

## Abstract

Symbiotic digestion of lignocellulose in the hindgut of higher termites is mediated by a diverse assemblage of bacteria and archaea. During a large-scale metagenomic study, we reconstructed 15 metagenome-assembled genomes (MAGs) of *Bathyarchaeia* that represent two distinct lineages in subgroup 6 (formerly MCG-6) unique to termite guts. One lineage (TB2; *Candidatus* Termitimicrobium) encodes all enzymes required for reductive acetogenesis from H_2_ and CO_2_ via an archaeal variant of the Wood**–**Ljungdahl pathway. This includes a novel 11-subunit hydrogenase, which possesses the genomic architecture of the respiratory Fpo-complex of other archaea but whose catalytic subunit is phylogenetically related to and shares the conserved [NiFe] cofactor-binding motif with [NiFe] hydrogenases of subgroup 4g. We propose that this novel Fpo-like hydrogenase provides the reduced ferredoxin required for CO_2_ reduction and is driven by the electrochemical membrane potential generated from the ATP conserved by substrate-level phosphorylation. Members of the other lineage (TB1; *Candidatus* Termiticorpusculum) are not capable of lithotrophic acetogenesis because they consistently lack hydrogenases and/or methylene-tetrahydromethanopterin reductase, a key enzyme of the pathway. Both lineages have the genomic capacity to reduce ferredoxin by oxidizing amino acids and might conduct methylotrophic acetogenesis using unidentified methylated compound(s). Our results indicate that *Bathyarchaeia* of subgroup 6 contribute to acetate formation in the guts of higher termites and substantiate the genomic evidence for reductive acetogenesis from organic substrates, including methylated compounds, in other uncultured representatives of the phylum.

## 1 Introduction

Although *Bathyarchaeia* are widespread in anoxic environments, their physiology is only poorly understood. In the absence of any isolates and only few microscopic observations of their cells (Collins *et al*., 2005, Kubo *et al*., 2012), our knowledge about this deep-branching lineage is based almost exclusively on amplicon libraries of archaeal 16S rRNA genes and metagenomic studies (reviewed by Zhou *et al*. (2018)).

Ribosomal RNA genes affiliated with the “Miscellaneous Crenarchaeotal Group” (MCG) had already been recovered in early analyses of archaeal diversity in diverse anoxic habitats (e.g., Schleper *et al*., 1997, Inagaki *et al*., 2003, Ochsenreiter *et al*., 2003), including the intestinal tract of termites (Friedrich *et al*., 2001). Meanwhile, an enormous diversity of sequences from this group, which comprises numerous deep-branching lineages, has been recovered from a wide range of marine and freshwater habitats and terrestrial environments (e.g., Kubo *et al*., 2012, Fillol *et al*., 2016). A few years ago, the MCG was elevated to the phylum level (*Bathyarchaeota*; Meng *et al*., 2014), but the most recent genome-based taxonomy demoted them again to the class level (*Bathyarchaeia*; Rinke *et al*., 2020). While the rank of the taxon is not relevant in the current context, we maintained the subgroup numbering used in previous studies (e.g., Kubo *et al*., 2012; Lazar *et al*., 2016) but replaced the prefix ‘MCG-’ with the prefix ‘Bathy-’ (Yu *et al*., 2018).

The abundance of *Bathyarchaeia* in many anoxic habitats implies potentially important roles in biogeochemical cycles (Evans *et al*., 2015; He *et al*., 2016). Reconstruction of metagenome-assembled genomes (MAGs) provided information concerning the metabolic capacities of *Bathyarchaeia* and inspired predictions of their putative roles in anoxic sediments (reviewed by Zhou *et al*., 2018). Several studies suggested that *Bathyarchaeia* are organotrophic and utilize a variety of organic substrates (e.g., Meng *et al*., 2014; He *et al*., 2016; Lazar *et al*., 2016). The discovery of genes encoding a methyl-coenzyme M reductase (Mcr) complex and a complete Wood–Ljungdahl pathway in bathyarchaeon BA1 provided the first evidence of methanogenesis outside the Euryarchaeota (Evans *et al*., 2015). Other studies detected key enzymes of the pathway in bathyarchaeal genomes of several subgroups and proposed that these lineages are involved in reductive acetogenesis from CO_2_ (He *et al*., 2016, Lazar *et al*., 2016).

Considering the putative roles of *Bathyarchaeia* in methanogenesis and reductive acetogenesis and the evidence for the utilization of lignin-derived methoxy groups (Yu *et al*., 2018), the presence of this group in termite guts is intriguing. Termites efficiently digest wood and other lignocellulosic substrates, either sound or in different stages of humification (Brune, 2014), in symbiosis with a specialized gut microbiota housed in their enlarged hindgut compartments (Brune and Dietrich, 2015). Hydrogen produced in microbial fermentation processes serves as electron donor for the reduction of CO_2_, yielding acetate and methane as major products (Breznak and Switzer, 1986; Brauman et al., 1992). Methanogenesis in termite guts involves a diverse assemblage of hydrogenotrophic and methyl-reducing archaea (Brune, 2018), but reductive acetogenesis, which can contribute up to two-thirds of total acetate production, has so far been considered a bacterial activity.

In lower termites, reductive acetogenesis has been attributed to homoacetogenic members of the phylum *Spirochaetes*) (e.g., Leadbetter *et al*., 1999, Ohkuma *et al*., 2015) and a novel lineage of uncultured *Deltaproteobacteria* (Rosenthal *et al*., 2013; Ikeda-Ohtsubo *et al*., 2016). In higher termites (family Termitidae), which diverged from the lower termites about 50 million years ago (Bucek *et al*., 2019), the situation is more complex. Particularly in the humus-feeding and soil-feeding groups, where the potential rates of reductive acetogenesis decrease in favor of methanogenesis (Brauman *et al*., 1992; Tholen and Brune, 1999), spirochetes are less abundant than in wood-feeding groups (Mikaelyan *et al*., 2016). A study based on the formyltetrahydrofolate synthetase (FTHFS) gene, a key enzyme of the Wood–Ljungdahl pathway that has been used as a marker for reductive acetogenesis, indicated that the community of potential acetogens shifts from spirochetes in lower termites to clostridia in higher termites (Ottesen and Leadbetter, 2011).

In a large-scale metagenomic study of the gut microbiota of eight higher termites, we obtained 15 metagenome-assembled genomes (MAGs) assigned to *Bathyarchaeia* (Hervé *et al*., 2020). Preliminary analysis revealed that they fell into a cluster comprising mainly termite gut MAGs, with members of Bathy-1 and Bathy-6 as next relatives. Here, we conducted detailed phylogenomic analyses of these MAGs and investigated their potential capacity for methanogenesis and reductive acetogenesis using a genome-centric approach.

## 2 Results and Discussion

### 2.1 Phylogeny of termite gut *Bathyarchaeia*

Bathyarchaeal MAGs were recovered from seven of the eight higher termites investigated, regardless of their feeding group (Hervé *et al*., 2020; Table 1). Their absence from *Microcerotermes parvus* is most likely caused by the low total number of MAGs obtained from the metagenomes of this species. Based on average nucleotide identity, the MAGs were assigned to nine phylotypes (Table 1). MAGs of the same phylotype were always derived from different gut compartments of the same host species, indicating that they most likely represent bathyarchaeal populations distributed along the entire hindgut. Eleven of the 15 MAGs fulfill the criteria for high-quality MAGs (>90% complete and <5% contamination; Bowers *et al*., 2017). Except for phylotype 5, each phylotype is represented by at least one high-quality MAG, which allows robust inference of metabolic potentials (Nelson *et al*., 2020).

**Table 1.**
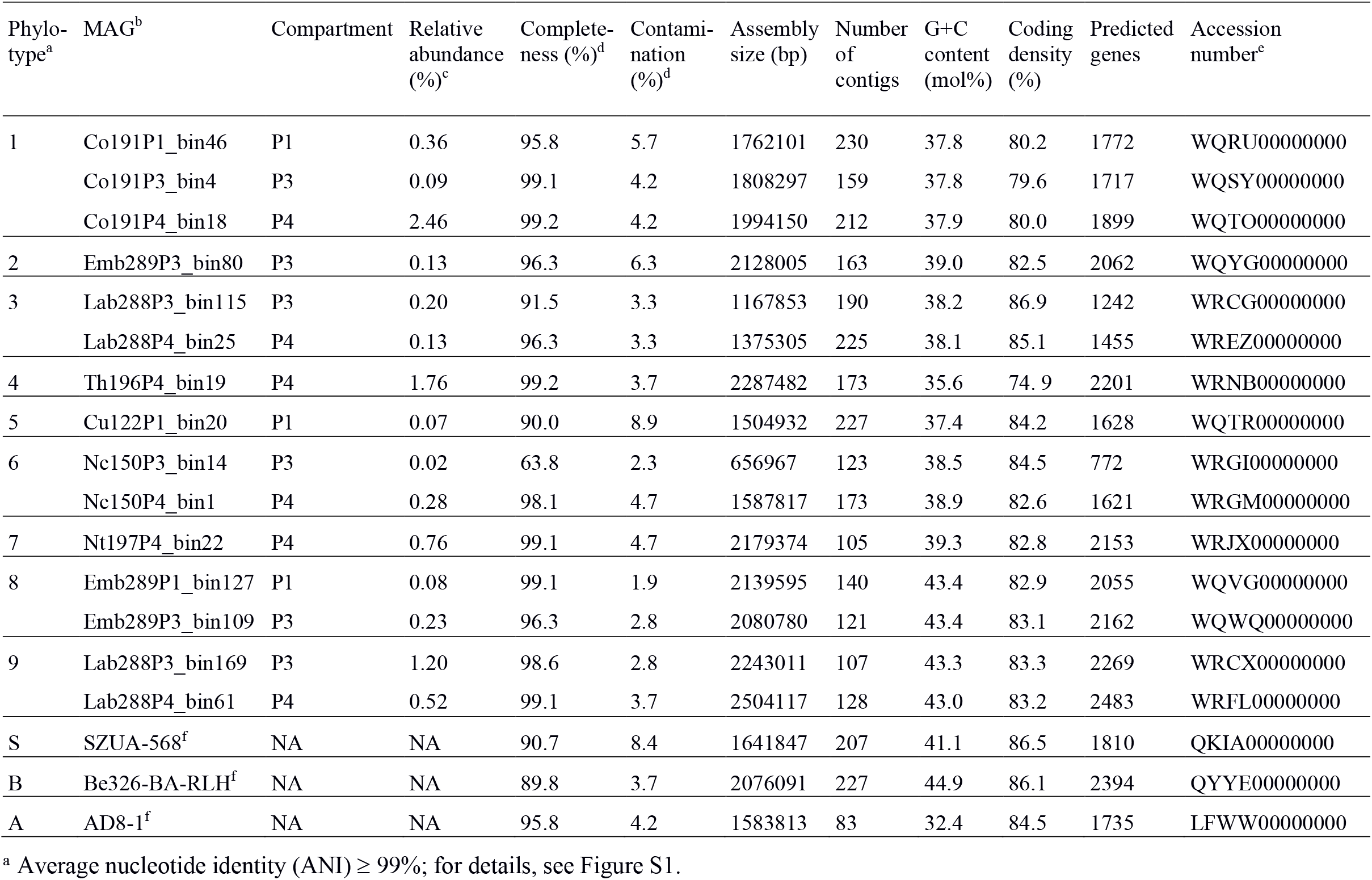
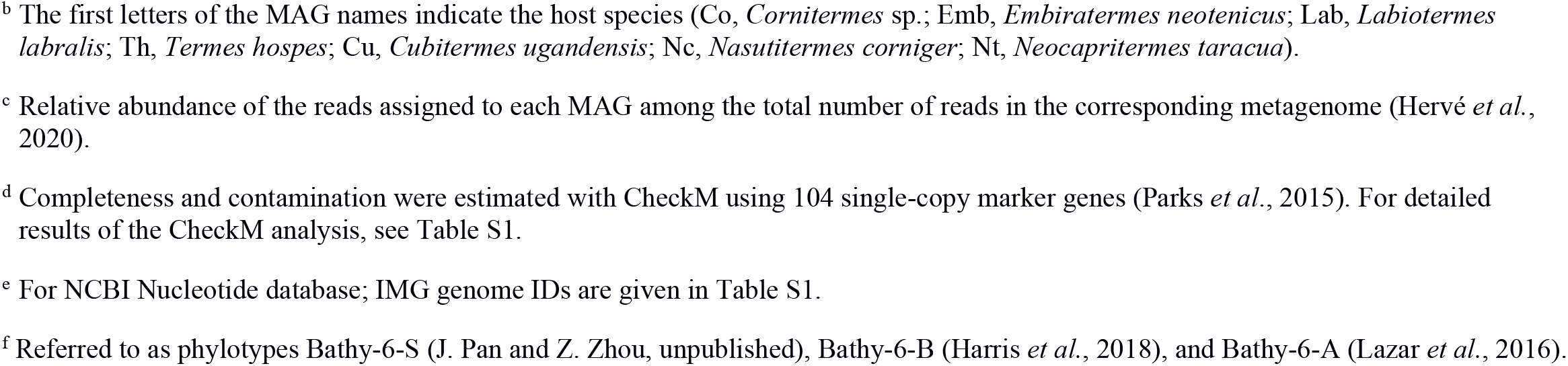
Characteristics of the MAGs of *Bathyarchaeia* from termite guts and other members of Bathy-6 included in the analyses.

| Phylo-<br>type<sup>a</sup> | MAG<sup>b</sup> | Compartment | Relative<br>abundance<br>(%)<sup>c</sup> | Complete-<br>ness (%)<sup>d</sup> | Contami-<br>nation<br>(%)<sup>d</sup> | Assembly<br>size (bp) | Number<br>of<br>contigs | G+C<br>content<br>(mol%) | Coding<br>density<br>(%) | Predicted<br>genes | Accession<br>number<sup>e</sup> |
| --- | --- | --- | --- | --- | --- | --- | --- | --- | --- | --- | --- |
| 1 | Co191P1_bin46 | P1 | 0.36 | 95.8 | 5.7 | 1762101 | 230 | 37.8 | 80.2 | 1772 | WQRU000000000 |
|  | Co191P3_bin4 | P3 | 0.09 | 99.1 | 4.2 | 1808297 | 159 | 37.8 | 79.6 | 1717 | WQSY000000000 |
|  | Co191P4_bin18 | P4 | 2.46 | 99.2 | 4.2 | 1994150 | 212 | 37.9 | 80.0 | 1899 | WQTO000000000 |
| 2 | Emb289P3_bin80 | P3 | 0.13 | 96.3 | 6.3 | 2128005 | 163 | 39.0 | 82.5 | 2062 | WQYG000000000 |
| 3 | Lab288P3_bin115 | P3 | 0.20 | 91.5 | 3.3 | 1167853 | 190 | 38.2 | 86.9 | 1242 | WRCG000000000 |
|  | Lab288P4_bin25 | P4 | 0.13 | 96.3 | 3.3 | 1375305 | 225 | 38.1 | 85.1 | 1455 | WREZ000000000 |
| 4 | Th196P4_bin19 | P4 | 1.76 | 99.2 | 3.7 | 2287482 | 173 | 35.6 | 74.9 | 2201 | WRNB000000000 |
| 5 | Cu122P1_bin20 | P1 | 0.07 | 90.0 | 8.9 | 1504932 | 227 | 37.4 | 84.2 | 1628 | WQTR000000000 |
| 6 | Nc150P3_bin14 | P3 | 0.02 | 63.8 | 2.3 | 656967 | 123 | 38.5 | 84.5 | 772 | WRGI000000000 |
|  | Nc150P4_bin1 | P4 | 0.28 | 98.1 | 4.7 | 1587817 | 173 | 38.9 | 82.6 | 1621 | WRGM000000000 |
| 7 | Nt197P4_bin22 | P4 | 0.76 | 99.1 | 4.7 | 2179374 | 105 | 39.3 | 82.8 | 2153 | WRJX000000000 |
| 8 | Emb289P1_bin127 | P1 | 0.08 | 99.1 | 1.9 | 2139595 | 140 | 43.4 | 82.9 | 2055 | WQVG000000000 |
|  | Emb289P3_bin109 | P3 | 0.23 | 96.3 | 2.8 | 2080780 | 121 | 43.4 | 83.1 | 2162 | WQWQ000000000 |
| 9 | Lab288P3_bin169 | P3 | 1.20 | 98.6 | 2.8 | 2243011 | 107 | 43.3 | 83.3 | 2269 | WRCX000000000 |
|  | Lab288P4_bin61 | P4 | 0.52 | 99.1 | 3.7 | 2504117 | 128 | 43.0 | 83.2 | 2483 | WRFL000000000 |
| S | SZUA-568<sup>f</sup> | NA | NA | 90.7 | 8.4 | 1641847 | 207 | 41.1 | 86.5 | 1810 | QKIA000000000 |
| B | Be326-BA-RLH<sup>f</sup> | NA | NA | 89.8 | 3.7 | 2076091 | 227 | 44.9 | 86.1 | 2394 | QYYE000000000 |
| A | AD8-1<sup>f</sup> | NA | NA | 95.8 | 4.2 | 1583813 | 83 | 32.4 | 84.5 | 1735 | LFWW000000000 |

<sup>a</sup> Average nucleotide identity (ANI) ≥ 99%; for details, see Figure S1.

Phylogenomic analysis placed all phylotypes from termite guts within subgroup Bathy-6, an apical lineage of *Bathyarchaeia* that is well represented mostly in 16S rRNA gene libraries (He *et al*., 2016) but comprises only a few MAGs from marine or estuarine sediments and the deep subsurface (Figure 1). The MAGs from termite guts form two distinct lineages, TB1 (phylotypes 1–7) and TB2 (phylotypes 8 and 9). TB2 is a sister group of bathyarchaeon SZUA-568 (hereafter denoted as Bathy-6-S), a MAG retrieved from marine hydrothermal vent sediments. Other MAGs in the radiation of Bathy-6 are bathyarchaea BE326-BA-RLH (hereafter denoted as Bathy-6-B) and AD8-1 (hereafter denoted as Bathy-6-A). They are all high-quality MAGs and were included in the subsequent analyses (Table 1). Only bathyarchaeon SG8-32-3 (previously assigned to Bathy-1) was omitted because the completeness of the assembly (50.4%; based on our CheckM analysis) was too low for a reliable assessment of its metabolic capacity.

**Figure 1.**
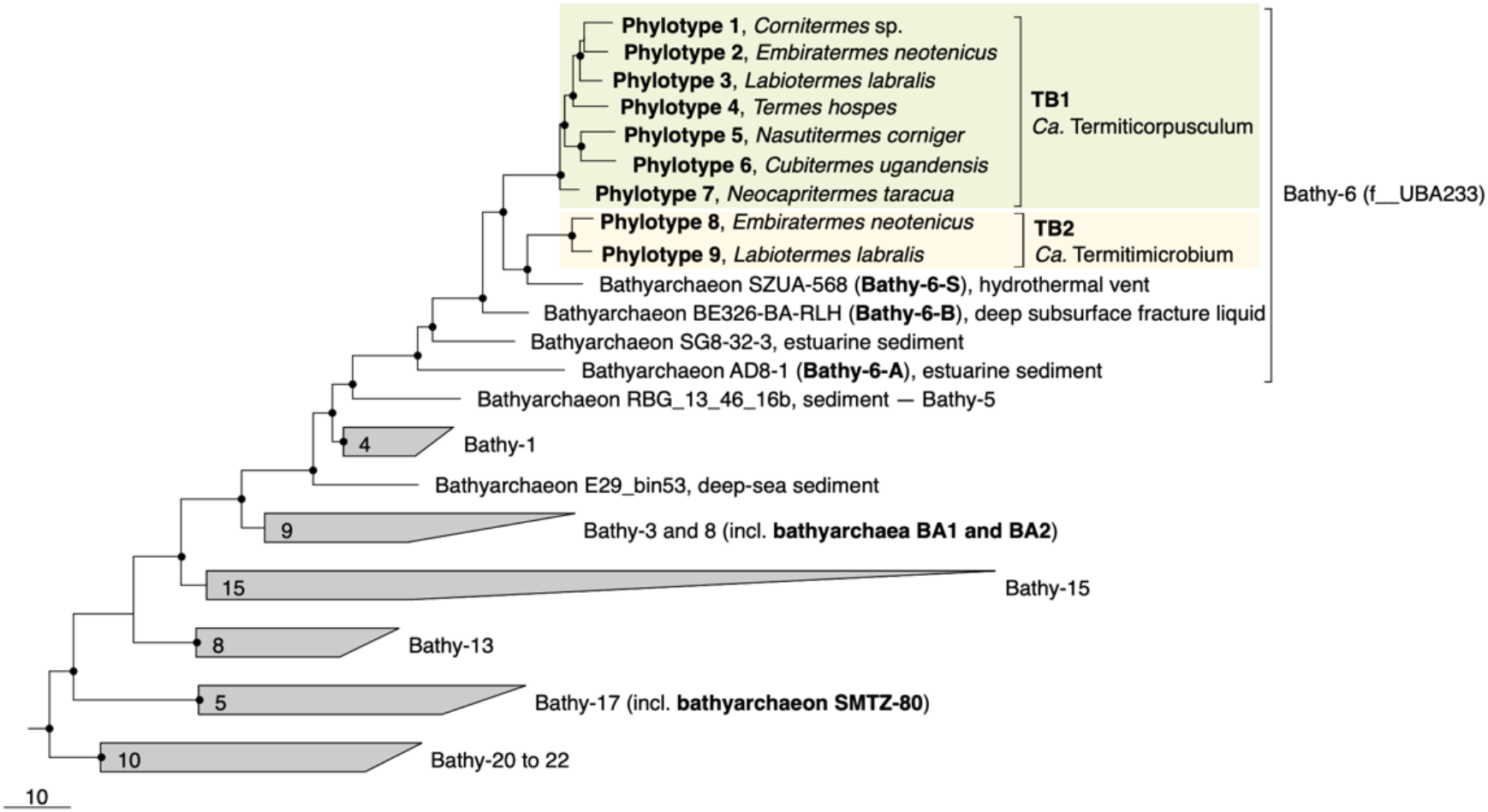
Genome-based phylogeny of termite gut *Bathyarchaeia*, illustrating the relationship of lineages TB1 and TB2 to other MAGs in the Bathy-6 subgroup (f UBA233 in the GTDB taxonomy). MAGs of other subgroups that are mentioned in the text are marked in bold. The maximum-likelihood tree was inferred from a concatenated alignment of 43 marker genes using the LG+F+I+G4 model and rooted with selected Crenarchaeota and Euryarchaeota as outgroup. A fully expanded tree with the accession numbers for all genomes is shown in the Supplementary Material (Supplementary Figure S2). The scale bar indicates 10 amino acid substitutions per site. Highly supported nodes (SH-aLRT, ● ≥ 95%, 1,000 replications) are indicated.

Predicted genome sizes (1.0 – 2.5 Mbp), G+C contents (37.4 – 43.4 mol%), and coding densities (74.9 – 86.9%) of the MAGs from termite guts are in the same range as those of the other representatives of this subgroup (Table 1). While the average nucleotide identity (ANI) values among the phylotypes of TB1 and TB2 ranges between 78.1 and 81.6%, the ANI values between members of TB1, TB2, and the other phylotypes of Bathy-6 are below the cut-off of the fastANI tool (< 75%; Supplementary Figure S1), indicating that each lineage represents a separate genus-level taxon. This is confirmed by the results obtained with the GTDB toolkit, which classified members of TB1 and TB2 as separate, genus-level lineages in the family UBA233 (order B26-1), a family that comprises also other members of Bathy-6. This indicates that TB1 and TB2 represent novel candidate genera in family UBA233, for which the names ‘*Candidatus* Termiticorpusculum’ and ‘*Candidatus* Termitimicrobium’ are proposed.

To identify the closest relatives of termite gut *Bathyarchaeia* and their respective habitats, we analyzed their phylogenetic position in the framework of rRNA genes available in public databases, which provides a much better coverage than the small number of MAGs of the Bathy-6 subgroup available to date (Figure 2). The 16S rRNA gene sequences encoded by the MAGs form a well-supported monophyletic group with all other sequences of *Bathyarchaeia* that were previously obtained from the hindguts of higher termites (Friedrich *et al*., 2001; Shi *et al*., 2015; Grieco *et al*., 2019). Although each ribotype appears to be specific for a particular host species, the internal topology of the termite clade is not well resolved due to the large number of short sequences and the absence of 16S rRNA genes from many MAGs. The sequences in the termite clade are most closely related to clones obtained from a manure pit (EU662668; J. Ding, unpublished) and an anaerobic digestor fed with vinasses (U81774; Godon et al., 1997), and fall into the radiation of bathyarchaeal lineages in freshwater sediments, salt marshes, and anaerobic wastewater bioreactors (group 1.3b; Ochsenreiter *et al*., 2003; Collins *et al*., 2005).

**Figure 2.**
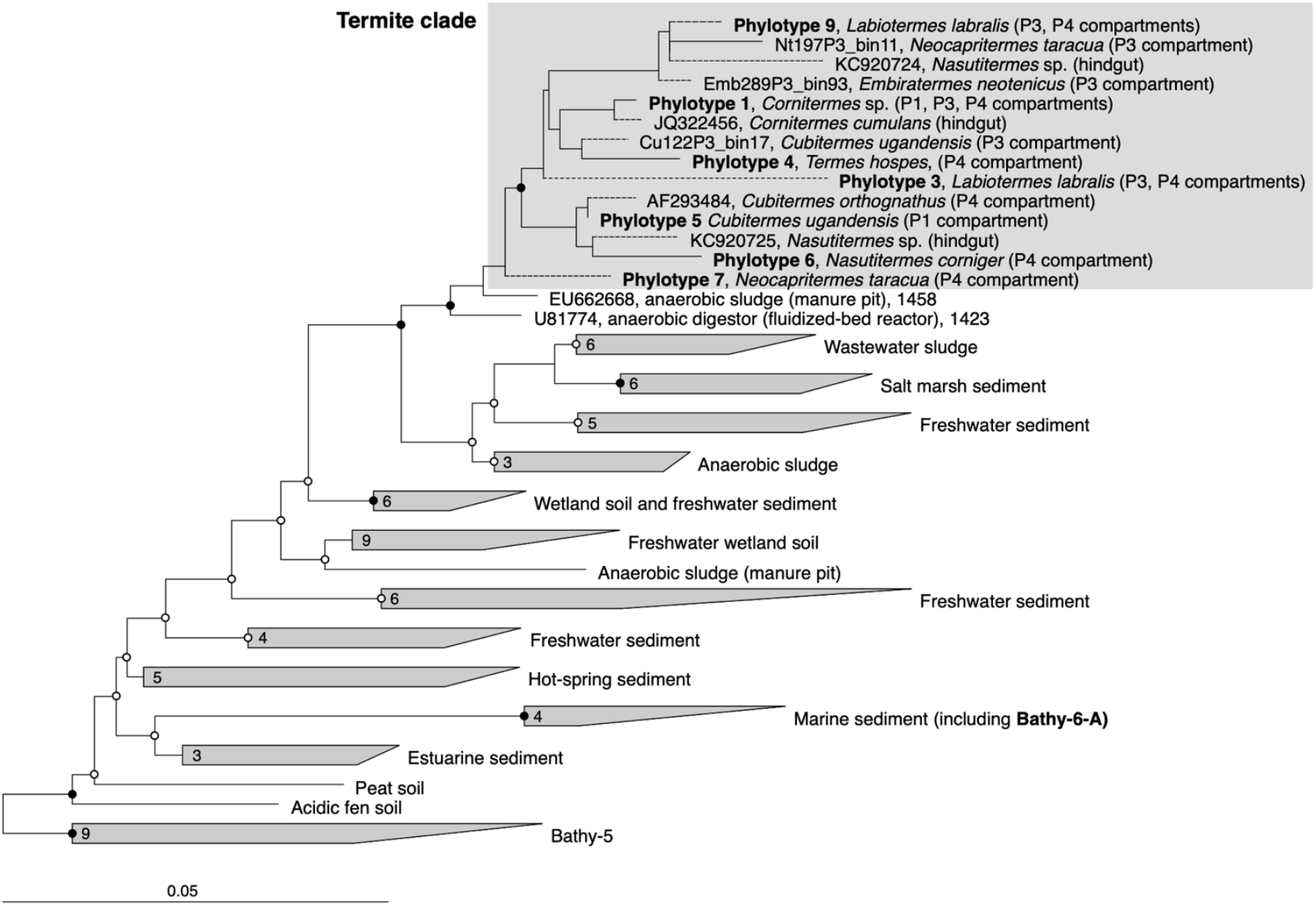
16S rRNA-based phylogeny of subgroup Bathy-6, indicating the placement of the termite clade among *Bathyarchaeia* from other environments. The maximum-likelihood tree is based on a curated alignment (1,424 positions) of all sequences in the SILVA database and their homologs retrieved from the bathyarchaeal MAGs and the low-quality bins obtained from the termite gut metagenomes (Hervé *et al*., 2020). The tree was rooted with members of Bathy-5 as outgroup. The scale bars indicate 0.05 nucleotide substitutions per site. SH-aLRT values (● ≥ 95%; ○ ≥ 80%, 1,000 replications) indicate node support. Branches marked with dashed lines indicate shorter sequences that were added using the parsimony tool. A fully expanded tree with the accession numbers of all sequences is shown in the Supplementary Material (Supplementary Figure S3).

### 2.2 Capacity for CO_2_-reductive acetogenesis

We investigated the presence of all genes required for methanogenesis and reductive acetogenesis in all members of Bathy-6 with sufficiently complete genomes (Figure 3). All members of TB2 (phylotypes 8 and 9) encode the complete set of genes required for the reduction of CO_2_ to acetyl-CoA via the archaeal version of the Wood–Ljungdahl pathway, using methanofuran (MFR) and tetrahydromethanopterin (H_4_MPT) as C_1_ carriers (Figure 4). Formyl-MFR dehydrogenase is molybdenum-dependent (FmdABCDF; Hochheimer *et al*., 1996) and not the tungsten-dependent paralog. A homolog of *fmdE*, which occurs in methanogens, was not found in any of the MAGs, which suggests that the absence of subunit E is a characteristic feature of the bathyarchaeal complex. It has been shown that the Fmd complexes of *Methanobacterium thermoautotrophicum* and *Methanosarcina barkeri* are active also without this subunit (Hochheimer *et al*., 1996, Vorholt *et al*., 1996). The CO dehydrogenase/acetyl-CoA synthase complex (CdhABCDE) and the (ADP-forming) acetyl-CoA synthetase (Acd; Musfeldt *et al*., 1999) are typical archaeal enzymes.

**Figure 3.**
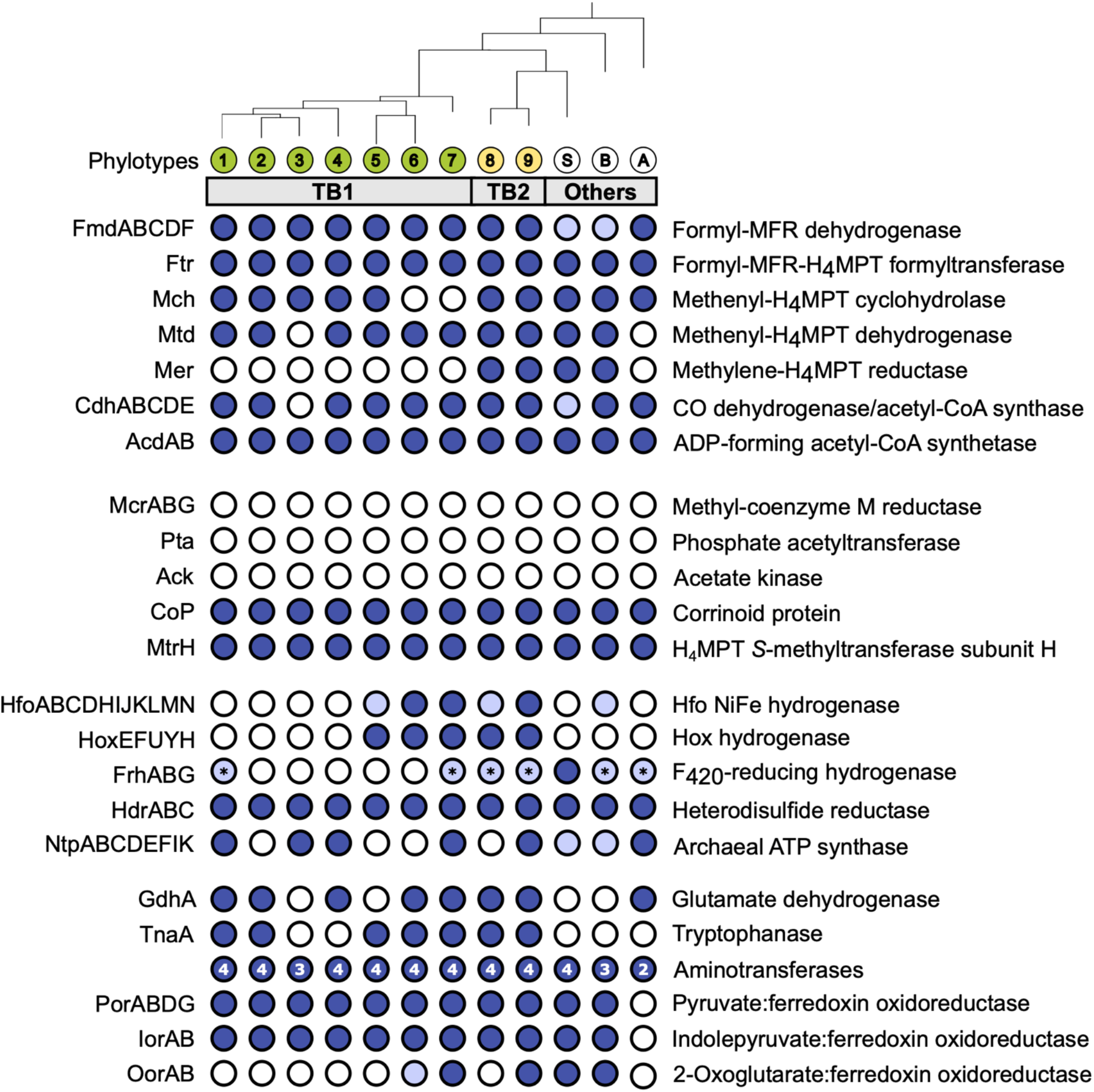
Gene functions encoded by termite gut bathyarchaea (TB1 and TB2) and other representatives of the Bathy-6 subgroup. All phylotypes with sufficiently complete genomes were included; their phylogenetic relationship was taken from Figure 1 (for strain designations, see Table 1). Colored circles indicate presence and open circles indicate absence of the respective function; light blue indicates that a gene set is incomplete. The number of aminotransferases encoded by each phylotype are indicated in the circle. The asterisk (*) in FrhABG indicates that only FrhB is present. If a phylotype is represented by more than one MAG, the annotation results were combined; details can be found in the Supplementary Material (Table S2). H_4_MPT: tetrahydromethanopterin, MFR: methanofuran, Fpo: F_420_:methanophenazine oxidoreductase.

**Figure 4.**
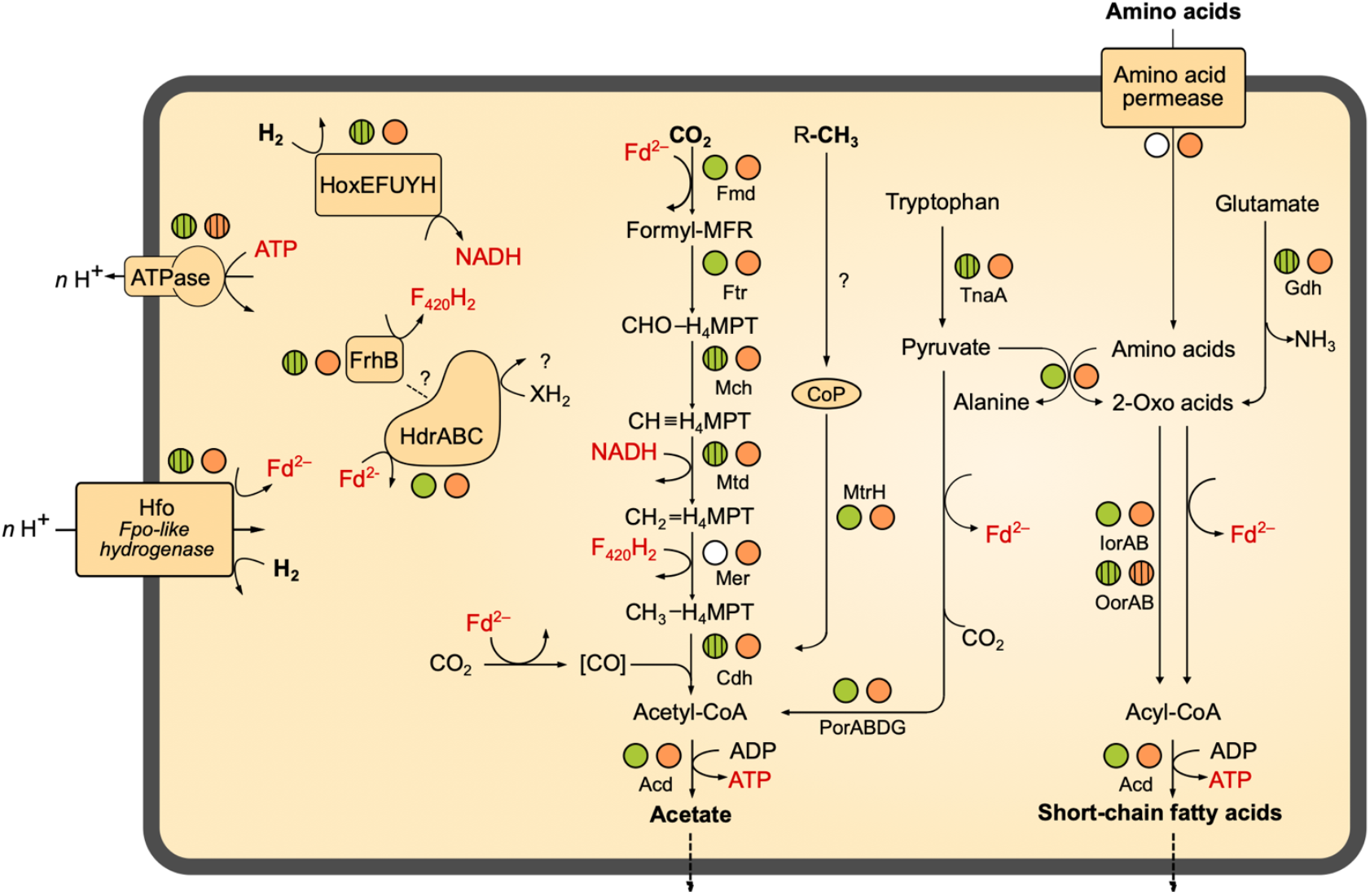
Metabolic map of termite gut *Bathyarchaeia*. The circles next to each enzyme indicate the presence of the coding genes in TB1 (green) and TB2 (orange), respectively. Striped circles indicate that a function is not encoded by all genomes of the respective group; a white filling indicates absence from all MAGs of the group (see also Figure 3). The directionality of Fpo-like hydrogenase (Hfo) and ATP (synth)ase is explained in the text. A detailed list of genes present in the respective MAGs is provided as Supplementary Material (Supplementary Table S2).

Enzymes characteristic for the bacterial Wood**–**Ljungdahl pathway (FTHFS, methylene-THF cyclohydrolase/dehydrogenase, and methylene-THF reductase), which had been identified in MAGs of Bathy-3, -8 and -17 (Evans *et al*., 2015, Zhou *et al*., 2018), were not encoded by any member of Bathy-6. Also, phosphate acetyltransferase and acetate kinase, which are responsible for substrate-level phosphorylation in fermenting bacteria, were absent from all MAGs.

The same gene sets as in TB2 are encoded also by the more basal Bathy-6-S and Bathy-6-B (Figure 3), which indicates that the capacity to produce acetate from CO_2_ might be a plesiomorphic trait of the Bathy-6 subgroup. The consistent absence of a key enzyme of the archaeal Wood**–**Ljungdahl pathway, methylene-H_4_MPT reductase (Mer), from all seven phylotypes (11 MAGs) of the TB1 lineage and from the most basal member of the subgroup, Bathy-6-A, suggests that the capacity to reduce CO_2_ to the methyl level was lost at least twice during the evolutionary radiation of Bathy-6.

Homologs of the methyl-coenzyme M reductase (Mcr) complex, which encodes the key enzyme of methanogenesis, were not detected in any of the MAGs. Our observation contrasts with the report of Harris *et al*. (2018), who claimed that Bathy-6-B might represent an anaerobic methane oxidizer. However, their conclusion is based on the recovery of a 265-bp gene fragment classified as an *mcrA* gene in the original metagenome from which Bathy-6-B was assembled, i.e., not from the metagenomic bin. Considering also that the gene fragment in question shows highest similarity to a homolog from an uncultured euryarchaeal methanogen (GenBank: JX907770.1), it seems safe to conclude that members of the Bathy-6 subgroup are not methanogenic.

Although the capacity of *Bathyarchaeia* for reductive acetogenesis from CO_2_ has been claimed repeatedly for several subgroups (He *et al*., 2016; Lazar *et al*., 2016; Zhou *et al*., 2018; Yu *et al*., 2018), the evidence was never fully conclusive. Actually, the recent, comprehensive survey of all bathyarchaeal MAGs compiled by Zhou *et al*. (2018) lists only two MAGs that encode all genes required to operate the entire Wood**–**Ljungdahl pathway. One is the putatively methanogenic BA1 (Bathy-8) from a deep aquifer (Evans *et al*., 2015); the other is bathyarchaeon ex4484_135 (Bathy-15) from marine hydrothermal sediment (Dombrowski *et al*., 2017).

### 2.3 Capacity for methylotrophic acetogenesis

Since all members of Bathy-6 encode a complete CO dehydrogenase/acetyl-CoA synthase (Cdh) complex (Figure 3), they might still synthesize acetyl-CoA using methyl groups derived from external sources. In all homoacetogenic bacteria and methylotrophic methanogens studied to date, the methyl transferase systems consist of three components: (i) a set of substrate-specific methyl transferases (MT-I), (ii) their cognate methyl-accepting corrinoid proteins (CoP), and (iii) a second methyl transferase (MT-II) that transfers the methyl group of methyl-CoPs to THF (bacteria) or coenzyme M (archaea) (van der Meijden *et al*.,1983; Kreft and Schink, 1994; Kremp *et al*., 2018; Supplementary Figure S4A). We found that all MAGs of Bathy-6 encode CoPs that fall into the radiation of homologs assigned to other uncultured Archaea, with the CoPs of the di- and trimethylamine-specific methyltransferase systems (MtbC and MttC) of *Methanomassiliicoccus luminyensis* (Kröninger *et al*., 2017) and *Acetobacterium woodii* (Kremp *et al*., 2018) as closest relatives with a reliable functional annotation (Supplementary Figure S4). However, unlike the situation in methylotrophic bacteria and euryarchaea, where the CoP gene is co-localized with the gene of the cognate substrate-specific MT-I homologs (MtbB or MttB), the CoP gene of Bathy-6 is flanked by a gene encoding subunit H of tetrahydromethanopterin *S*-methyltransferase (MtrH; Supplementary Figure S4B).

In many methanogenic archaea, MtrH is part of the energy-conserving MtrA–H complex and catalyzes the transfer of the (CO_2_-derived) methyl group from methyl-tetrahydromethanopterin to the corrinoid prosthetic group of MtrA (Hippler and Thauer, 1999). However, in obligately methyl-reducing methanogens (Galagan *et al*., 2002; Borrel *et al*., 2014; Lang *et al*., 2015), which methylate CoM via their diverse methyltransferase systems (see above), the Mtr complex is absent. The presence of an isolated *mtrH* gene colocalized with a CoP gene has been observed also in the putatively methanogenic BA1 and BA2 (*Bathyarchaeia*) and several MAGs related to ‘*Ca*. Methanomethylicus mesodigestum’ (*Thermoproteota*). It was proposed that the encoded proteins represent methyltransferase systems, which prompted the hypothesis that these uncultured lineages are methylotrophic methanogens (Evans *et al*. 2015; Vanwonterghem *et al*. 2016).

It is tempting to assume that also the CoP–MtrH couple of Bathy-6 is involved in the transfer of methyl groups from so-far unidentified, substrate-specific methyltransferases to H_4_MPT (Figure 4). However, a catabolic role of the CoP–MtrH couple is not the only possible interpretation. In ‘*Ca*.

Methanomethylicus mesodigestum’, the genes are colocalized with a homolog of *metE* encoding methionine synthase (Supplementary Figure S4B), it is also possible that the CoP–MtrH couple of Bathy-6 is involved in anabolic reactions that transfer methyl groups (provided by the cleavage of acetyl-CoA) from H_4_MPT to an unknown acceptor.

### 2.4 Hydrogen as electron donor

The operation of the Wood**–**Ljungdahl pathway requires electron donors in the form of reduced ferredoxin, NADH, and in the case of archaea, also reduced cofactor F_420_ (F_420_H_2_). The reduction of ferredoxin with H_2_ is a critical step because it is endergonic at low hydrogen partial pressures and requires either an energy-converting hydrogenase or a flavin-based electron bifurcation system (Thauer *et al*., 2008, Buckel and Thauer, 2013).

Hydrogenases are present only in TB2 and the basal lineages of TB1 (Figure 3). One is a cytosolic, bidirectional [NiFe] hydrogenase of Subgroup 3d, which use NAD as electron acceptor (Greening *et al*., 2016). Phylogenetic analysis of the gene encoding the large subunit (*hoxH*) placed all homologs in a sister position to the Hox hydrogenases of phototrophic bacteria (Supplementary Figure S5). The gene order in the *hoxEFUYH* cluster is the same as in the gene clusters of other Hox complexes, which encode a prototypical heterodimeric [NiFe]-hydrogenase moiety (HoxHY) and a diaphorase moiety (HoxEFU); HoxEFU is homologous to the NuoEFG module of complex I and mediates the electron transport to NAD(P) (Eckert et al., 2012). Although members of Group 3 are called “bidirectional hydrogenases”, hydrogen formation requires reduced ferredoxin or flavodoxin as electron donor (Gutekunst et al. 2014).

All MAGs that encode a Hox hydrogenase also possess a gene cluster that resembles that encoding the F_420_:methanophenazine oxidoreductases (Fpo) of Euryarchaeota and the NADH:quinone oxidoreductases (Nuo) of Bacteria (complex I) (Figure 5). As in other basal lineages of complex I homologs, the FpoFO and NuoEFG modules, which provide substrate specificity for F_420_H_2_ or NADH, respectively, are absent (Moparthi and Hägerhäll, 2011).

**Figure 5.**
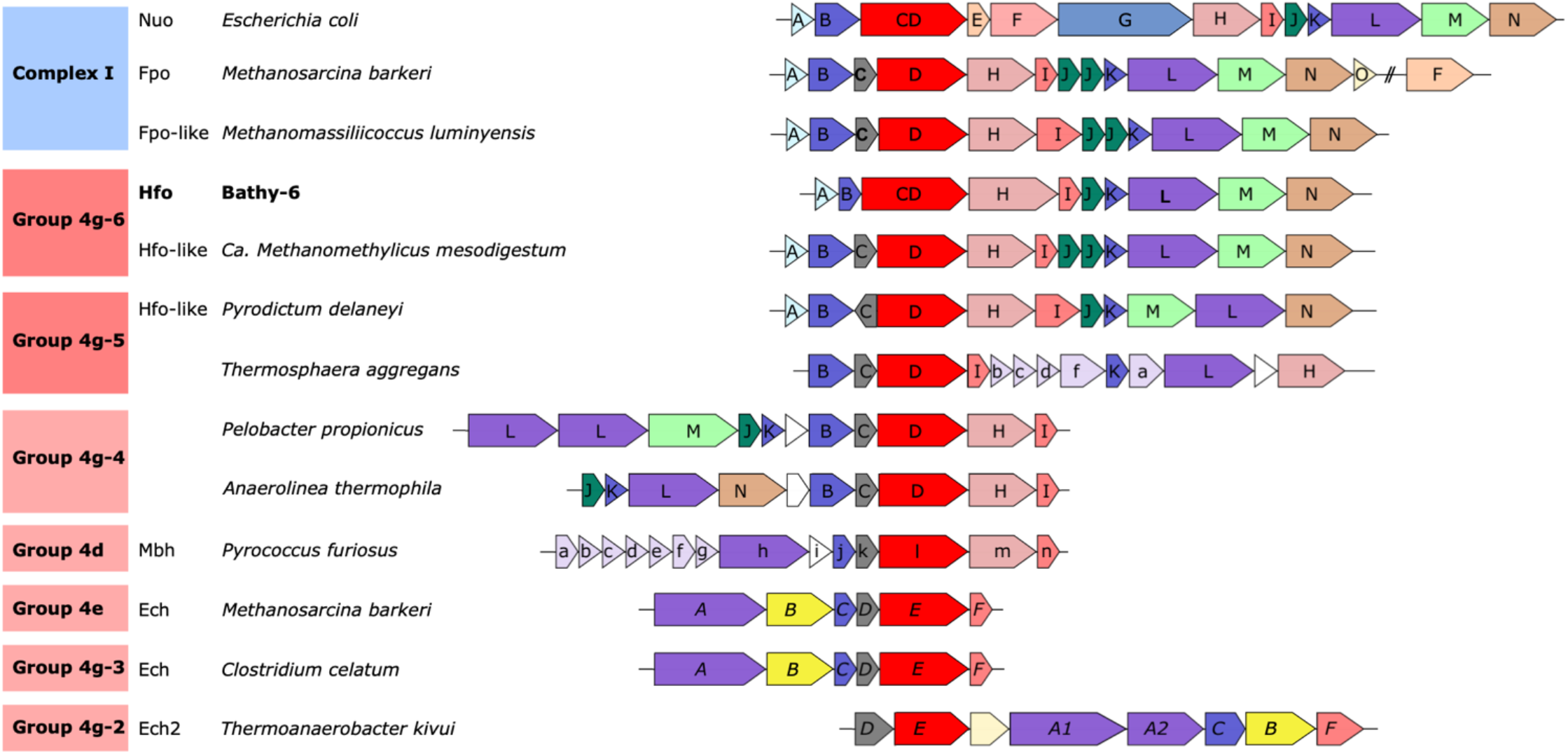
Genomic architecture of the gene clusters encoding the respiratory complex I (Nuo and Fpo) and the ancestral [NiFe] hydrogenases in Group 4. Colors indicate homologous genes; The phylogenetic analysis of the catalytic subunit of [NiFe] hydrogenases and its homologs (labeled in red) is shown in Figure 6. The font style of the gene labels indicates differences in the subunit nomenclature of Nuo/Fpo (uppercase), Mbh (lowercase), and Ech (italics).

However, six of the 11 subunits common to the Fpo and Nuo complexes are homologous to subunits of the energy-converting [NiFe] hydrogenases of Group 4, which are ancestral to the respiratory complex I (Friedrich and Scheide, 2000). Classification with HydDB placed the D subunit of the 11-subunit complex of the Bathy-6 MAGs among the catalytic subunits of [NiFe] hydrogenases in Subgroup 4g. The hydrogenases in Subgroup 4g are not only structurally heterogeneous and differ fundamentally both in the number of their subunits and the arrangement of their coding genes (Greening *et al*., 2016; Schoelmerich and Müller, 2019; Figure 5), but their large subunits also fall into separate phylogenetic lineages (Groups 4g-1 to 4g-6; Figure 6). The genomic architecture of the Fpo-like hydrogenase complex of Bathy-6 (hereafter referred to as Hfo) closely resembles that of *Ca*. Methanomethylicus mesodigestum (*Thermoproteota*) and *Pyrodictium delaney* (*Crenarchaeota*) (Figure 5), whose large subunits represent phylogenetic sister groups (4g-5 and 4g-6) that are distinct from the other lineages (Figure 6). Interestingly, the complex of *Thermosphaera aggregans* (Subgroup 4g-5) and other members of *Desulfurococcales* (not shown) seems to deviate from the Hfo-like structure and contains homologs of the Mbh complex of *Pyrococcus furiosus* (Figure 5).

**Figure 6.**
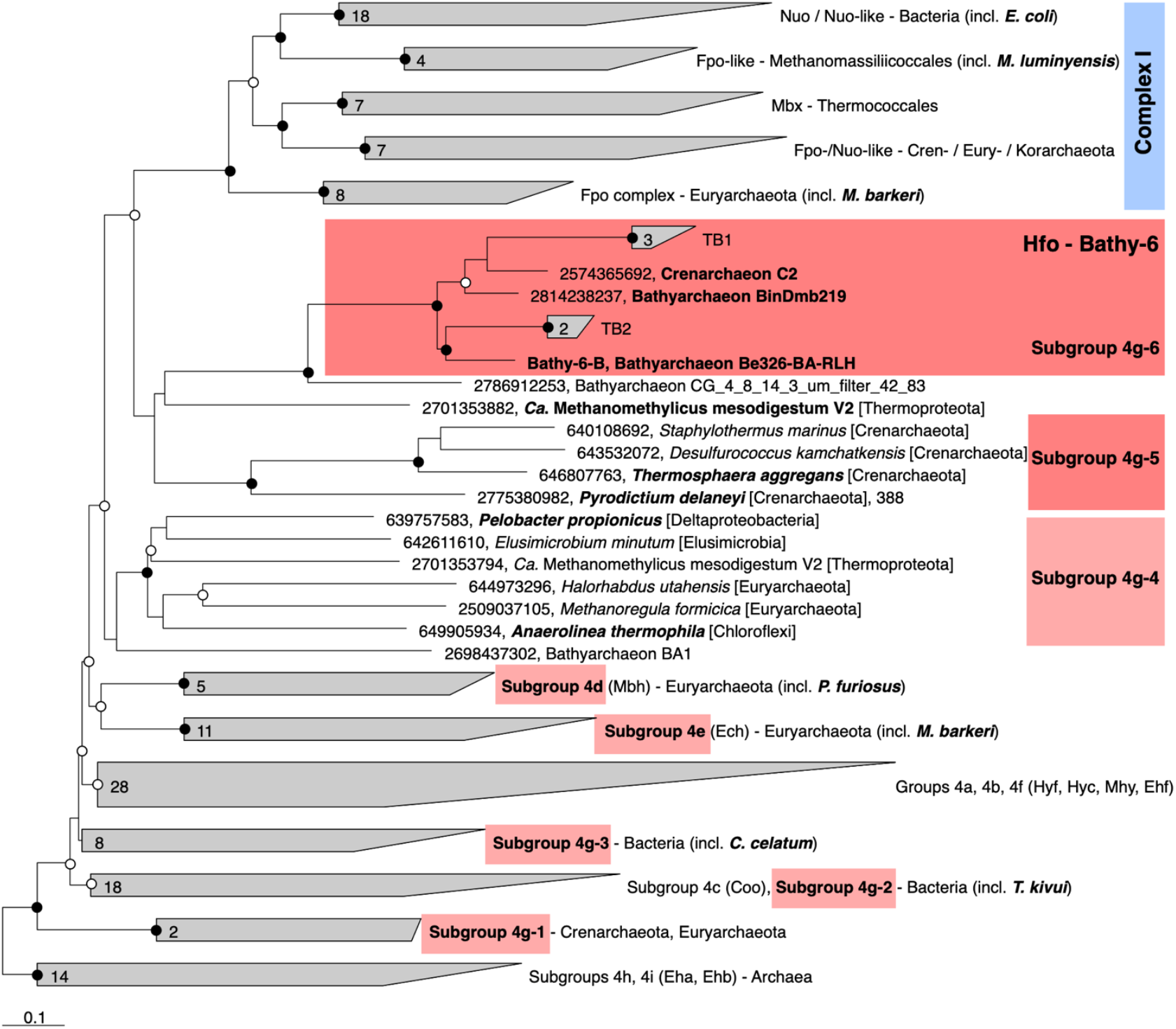
Phylogeny of the catalytic subunits of Group 4 [NiFe] hydrogenases and their homologs in respiratory complex I (FpoD and NuoD). The maximum-likelihood tree is based on a curated alignment of the deduced amino acid sequences; the scale bar indicates 0.1 amino acid substitutions per site. SH-aLRT values (● ≥ 95%; ○ ≥ 80%, 1,000 replications) indicate node support. The genomic context of the highlighted genes is shown in Figure 5. Gene numbers indicate IMG/Mer gene IDs.

None of the hydrogenases of Subgroup 4g have been biochemically characterized, but they are presumed to couple the formation of H_2_ from reduced ferredoxin to the formation of an electrochemical membrane potential (Greening *et al*., 2016; Søndergaard et al. 2016; Schoelmerich and Müller, 2019). This is in agreement with biochemical data obtained for the Fpo-like 11-subunit complex of methanogenic *Euryarchaeota*, which generate an electrochemical membrane potential during electron transport from reduced ferredoxin to methanophenazine (*Methanosaeta*; Welte and Deppenmeier, 2011) or a so far unidentified electron acceptor (*Methanomassiliicoccales*; Kröninger *et al*., 2016).

The presence of an Fpo-like 11-subunit hydrogenase in *Bathyarchaeia* is most interesting from an evolutionary perspective, since it represents the first [NiFe] hydrogenase with the genomic architecture of complex I. The coordination sites of the [NiFe] cofactor on the large subunit of all [NiFe] hydrogenases (L1 and L2 motifs; Vignais and Billoud, 2007), which are no longer conserved in NuoD and FpoD, are present in all Bathy-6 homologs (Figure 7). This adds to the evidence that the 11-subunit complex of *Bathyarchaeia* is not a respiratory complex but is instead a novel energy-converting hydrogenase that catalyzes the reduction of ferredoxin with H_2_ using the electrochemical membrane potential (Figure 4).

**Figure 7.**
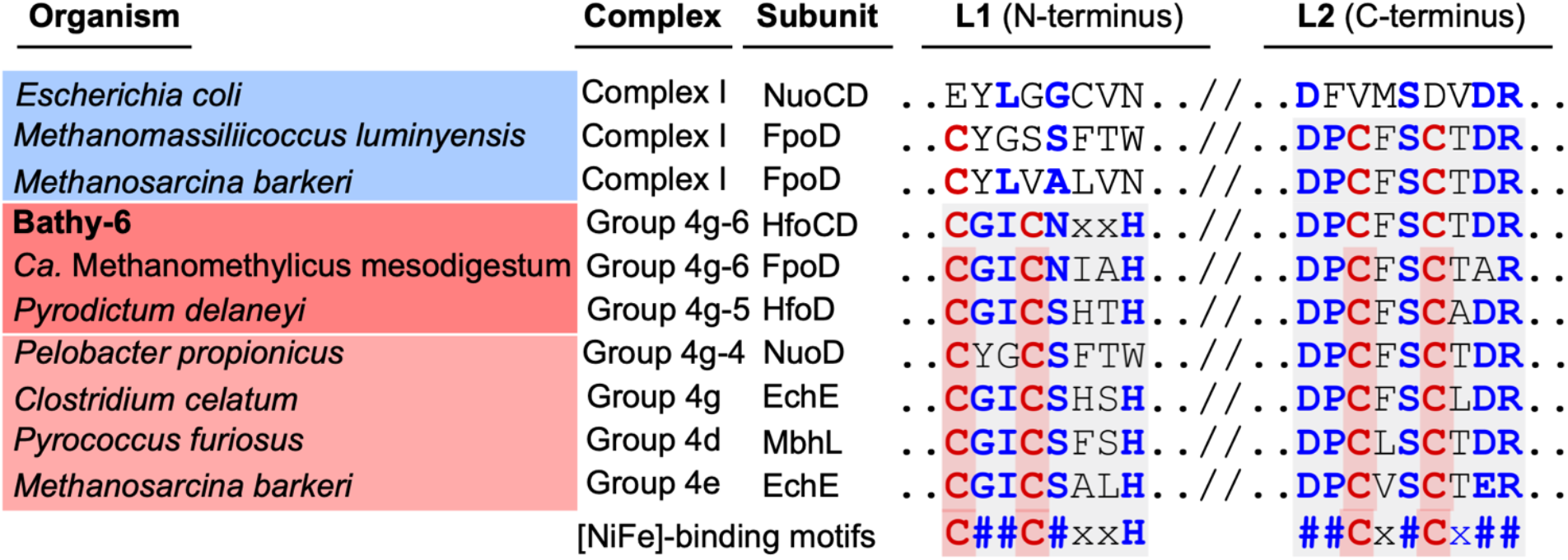
Comparison of the [NiFe]-binding motifs (L1 and L2) in the large subunits of selected Group 4 [NiFe] hydrogenases with the corresponding amino acid residues (IUPAC code) of their homologs in the respiratory complex I (Nuo and Fpo). Gray shading indicates that the typical motifs of [NiFe] hydrogenases are present (L1 motif: C[GS][ILV]C[AGNS]xxH; L2 motif: [DE][PL]Cx[AGST]Cx[DE][RL]; Vignais and Billoud, 2007). The four cysteine residues that coordinate the [NiFe] cluster are in red; other conserved residues are in blue.

While Hox and Fpo-like hydrogenase in TB2 should provide the NADH and reduced ferredoxin required to operate the Wood**–**Ljungdahl pathway, the source of F_420_H_2_ as potential electron donor for methylene-H_4_MPT reductase (Mer) remains unclear. A complete gene set encoding F_420_-reducing [NiFe] hydrogenase (FrhABG, subgroup 3a; Supplementary Figure S5) is present only in Bathy-6-A. All members of TB2 and several phylotypes of TB1 encode a homolog of FrhB, an iron–sulfur flavoprotein with an F_420_-binding site, but not the hydrogenase subunits (Figure 3). It is possible that FrhB is involved in the reduction of F_420_ via an interaction with HdrABC, as proposed for the methane-oxidizing *Ca*. Methanoperedens spp. (Arshad *et al*., 2015).

The only member of subgroup Bathy-6 that encodes a complete FrhABG is Bathy-6-A. It is also the only MAG that encodes a methylviologen-dependent [NiFe] hydrogenase (MvhADG; Supplementary Figure S5, subgroup 3c), which forms an electron-bifurcating complex with the soluble heterodisulfide reductase (HdrABC) and catalyzes the hydrogen-dependent reduction of ferredoxin and the heterodisulfide of coenzyme M (CoM) and coenzyme B (CoB) in methanogens (Kaster *et al*., 2011). The presence of genes encoding HdrABC, MvhADG, and a complete Wood**–**Ljungdahl pathway in the putatively methanogenic BA1 (Bathy-3) provides strong evidence that BA1 is capable of hydrogenotrophic methanogenesis (Evans *et al*., 2015). In Bathy-6-A, however, the pathway is incomplete and the identity of the heterodisulfide reduced by Hdr remains unclear. Interestingly, the same constellation as in Bathy-6 has been recently reported for the bathyarchaeal MAG CR_14 from marine sediments, which represents another, novel subgroup of *Bathyarchaeia* (Farag *et al*., 2020).

### 2.5 Organic substances as electron donors

Most members of TB1 and all basal lineages of Bathy-6 lack Hox and Fpo-like hydrogenase (Figure 3), which means that they cannot grow lithotrophically with H_2_ as electron donor. However, the reduced Fd required to operate acetogenesis, either via the Wood**–**Ljungdahl pathway (TB2) or by methylotrophy (all phylotypes), could be provided also by the oxidation of organic substrates (Figure 4). Such organotrophic acetogenesis is common among bacterial homoacetogens (Schink, 1994; Drake, 1994). All Bathy-6 genomes (except Bathy-6-A) encode pyruvate:ferredoxin oxidoreductase (Por) and indolepyruvate:ferredoxin oxidoreductase (Ior), and some also encode 2-oxoglutarate:ferredoxin oxidoreductase (Oor), all of which catalyze the oxidative decarboxylation of 2-oxo acids to their corresponding acyl-CoA esters (Figure 3). The 2-oxo-acids would result from the transamination of amino acids via numerous aminotransferases encoded by all genomes; a putative amino acid permease, however is encoded only in TB2. ATP would be formed via the ADP-dependent acetyl-CoA synthetase, which accepts also other acyl substrates in *Pyrococcus furiosus* (Mai and Adams, 1996). Such pathways have been shown to operate in other archaea (*Pyrococcus furiosus, Thermococcus* spp.; Kengen and Stams, 1993, Heider *et al*., 1996) and in the insect gut-associated bacterium *Elusimicrobium minutum* (Herlemann, *et al*., 2009) during growth on glucose, where they result in a net formation of alanine.

The data compiled by Zhou *et al*. (2018) suggest that several lineages of *Bathyarchaeia*, including Bathy-6-A; Lazar *et al*., 2016), have the capacity to ferment various organic carbon compounds. However, genes encoding extracellular peptidases, which are numerous in other *Bathyarchaeia*, seem to be less prevalent in the MAGs of Bathy-6 and Bathy-1 (Feng *et al*., 2019), which suggests that members of these subgroups are limited to the utilization of amino acids or oligopeptides that are small enough to be transported across the cytoplasmic membrane.

A capacity of members of Bathy-6 to utilize sugars is not as apparent. Like Bathy-6-A (Lazar *et al*., 2016), all MAGs of TB1 and TB2 encode many genes of the classical Embden-Meyerhof-Parnas (EMP) pathway, including glyceraldehyde-3-phosphate dehydrogenase and phosphoglycerate kinase. However, all MAGs lack hexokinase, the alternative archaeal glycolytic enzymes (Bräsen et al., 2014), and most MAGs lack phosphofructokinase and pyruvate kinase. It is possible that EMP pathway functions only in gluconeogenesis; fructose bisphosphatase is present in all MAGs. Sugar transporters were not detected; the role of the lipooligosaccharide ABC transporter encoded by almost all phylotypes from termite guts (except phylotype 9) is not clear (Supplementary Table S3). The identification of a cellulolytic system in Bathy-6-A (Lazar et al., 2016) requires verification.

### 2.6 Energy conservation

In acetogenic bacteria growing on hydrogen and CO_2_, all ATP synthesized by substrate-level phosphorylation is consumed in the activation of formate. Therefore, energy conservation involves electron-transport phosphorylation, which is driven by the oxidation of reduced ferredoxin via membrane-bound electron-transport complexes (Schuchmann and Müller, 2014; Basen and Müller, 2017). By contrast, the activation of formate (i.e., the formation of formylmethanofuran) in the archaeal variant of the Wood**–**Ljungdahl pathway is not ATP-dependent but is instead driven by the reducing power of ferredoxin, yielding a full ATP per acetate produced via substrate-level phosphorylation. However, thermodynamics dictates that a fraction of this ATP has to be reinvested, because a metabolism where the net ATP yield exceeds the free-energy change of the reaction would become endergonic (Thauer et al., 2008).

Fermenting bacteria that lack respiratory chains energize their membrane by operating their ATP synthase in the reverse direction (Buckel and Thauer, 2013). Likewise, members of Bathy-6 that have the capacity to grow lithotrophically on hydrogen and CO_2_ (i.e., the phylotypes in TB2) might use part of the ATP gained by substrate-level phosphorylation to generate an electrochemical membrane potential that drives the reduction of ferredoxin via the Fpo-like hydrogenase complex (see above). Other energy-converting complexes that would allow generation of reduced ferredoxin, such as the Group-4 [NiFe] hydrogenases in acetogenic bacteria and methanogenic archaea (Ech, Künkel *et al*., 2001; Eha and Ehb, Tersteegen and Hedderich, 2001) or an NADH:Fd oxidoreductase complex (RnfABCDEG, Westphal *et al*., 2018) were not detected in any member of Bathy-6.

During fermentative growth, the Fpo-like hydrogenase complex (if present) might operate in the reverse direction, using reduced ferredoxin provided by the oxidation of organic substrates to produce H_2_ and generate an electrochemical membrane potential, like many fermenting bacteria with an energy-converting hydrogenase. In that context, it is intriguing that several phylotypes of TB1 and TB2 (Figure 3), and also bathyarchaeal MAGs from other subgroups (Evans *et al*., 2015; Zhou *et al*., 2018), do not encode an ATP synthase (neither the genes for the archaeal V-type ATP synthase nor for the bacterial equivalent were detected). If one disregards the possibility of incomplete genome assemblies, these organisms must generate their membrane potential (vital for any organism) by other means.

In principle, the Wood**–**Ljungdahl pathway is reversible and can also oxidize acetate to H_2_ and CO_2_ given the appropriate thermodynamic framework. This has been demonstrated in syntrophic cultures of “Reversibacter”-like microorganisms with hydrogenotrophic partners (Lee and Zinder, 1988; Schnürer et al. 1997) and has been suggested to occur also in *Bathyarchaeia* (Evans *et al*., 2015; Xiang *et al*., 2017). However, at least in the termite hindgut, where the hydrogen partial pressure is much higher than in sediments (Ebert and Brune, 1997; Schmitt-Wagner et al., 1999) and reductive acetogenesis often prevails over methanogenesis as electron sink (Brauman et al., 1992; Tholen et al. 1999, Tholen and Brune, 2000), an anaerobic oxidation of acetate is an unlikely scenario.

### 2.7 Ecological aspects

Although the proportion of archaeal rRNA in termite hindguts is relatively small (0.9–2.3% of all prokaryotic rRNA; Brauman *et al*., 2001), methanogenesis represents a substantial hydrogen sink (Brune, 2019). Considering that the proportion of reads assigned to bathyarchaeal MAGs in the hindgut metagenomes of higher termites (0.03–2.5%; avg. 0.69%) is four times higher than that assigned to euryarchaeal MAGs (0.02–0.79%; avg. 0.16%; Table S2 in Hervé *et al*., 2020), the population sizes of *Bathyarchaeia* might be sufficient to contribute significantly to acetogenesis, particularly in soil-feeding species.

However, the substrates of termite gut *Bathyarchaeia* remain open to speculation. While only members of TB2 have the genomic capacity for lithotrophic acetogenesis, almost all members of Bathy-6 have the capacity to ferment amino acids and might employ organotrophic acetogenesis from methylated substrates as an electron sink. Stable-isotope probing of salt marsh sediments indicated that members of Bathy-8 and Bathy-6 assimilate organic substrates, notably excluding proteins and inorganic carbon (Seyler *et al*., 2014). Yu *et al*. (2018), however, reported that addition of lignin to an estuarine sediment sample selectively stimulated the growth of Bathy-8 and the incorporation of carbon from ^13^C-bicarbonate into archaeal tetraether lipids, which suggests that members of Bathy-8 are methylotrophs that use lignin-derived methyl groups. Together with the potential capacity for methyl group utilization in many bathyarchaeotal MAGs (Seyler *et al*., 2014; Yu *et al*., 2018; this study), these results explain the observations of Lever *et al*. (2010), who found that porewater acetate in deep-subseafloor sediments was depleted in ^13^C relative to sedimentary organic matter and postulated that a substantial fraction of the acetate produced in marine sediments might stem from reductive acetogenesis, fueled by microbial fermentation products, molecular hydrogen, and the methoxy groups of lignin monomers.

The utilization of the methoxy groups of lignin-derived aromatic compounds is a common trait of many acetogenic bacteria (Schink *et al*., 1992, Drake, 1994). Methoxylated aromatic compounds are demethylated by the hindgut microbiota of termites (Brune *et al*., 1995), but the organisms responsible for this activity have not been identified. It is tempting to speculate that termite gut *Bathyarchaeia* are organoheterotrophic (TB1) or lithoheterotrophic (TB2) acetogens that utilize methylated compounds such as lignin derivatives as methyl group donors and reduce CO_2_ either with molecular hydrogen or with reducing equivalents derived from the oxidation of organic substrates.

It has been speculated that acetogenic archaea might have an energetic advantage over acetogenic bacteria, as they do not have to invest ATP to activate formate (He *et al*. 2016). However, the net synthesis of ATP is limited by the free-energy change of an acetogenic metabolism, which is independent of its reaction path and requires part of the ATP gained by substrate-level phosphorylation to be reinvested (e.g., for ferredoxin reduction; see above). Rather, it is feasible that the capacity for methylotrophic acetogenesis, which is less sensitive to low hydrogen partial pressures than hydrogenotrophic acetogenesis, provides an energetic advantage, analogous to the situation in methyl-reducing methanogens (Feldwert *et al*., 2020). Moreover, it has been argued that long generation times contribute to the difficulties surrounding the enrichment and isolation of *Bathyarchaeia* in the laboratory (Yu *et al*. 2018). In view of the relatively short residence time of organic matter in termite guts (24 – 48 h; Kovoor, 1967; Bignell *et al*., 1980), the growth rates of termite gut *Bathyarchaeia* must be high enough to avoid washout – unless they are attached to the intestinal surface.

### 2.8 Taxonomy

#### *Candidatus* Termiticorpusculum

##### Etymology

L. n. *termes -itis*, a worm that eats wood, a termite; ; L. neut. n. *corpusculum*, a little body, a particle; N.L. neut. n. *Termiticorpusculum*, a little body associated with termites Uncultured. Unclassified genus-level lineage in the Bathy-6 subgroup of *Bathyarchaeia* (Fig. 1; TB1 lineage). Comprises the phylotypes 1–7 (Table 1).

##### Habitat

The hindgut of higher termites

#### *Candidatus* Termitimicrobium

##### Etymology

L. n. *termes -itis*, a worm that eats wood, a termite; N.L. neut. n. *microbium*, microbe; from Gr. masc. adj. *mikros*, small; from Gr. masc. n. *bios*, life; N.L. neut. n. *Termitimicrobium*, small life(-form) associated with termites Uncultured. Unclassified genus-level lineage in the Bathy-6 subgroup of *Bathyarchaeia* (Fig. 1; TB2 lineage). Comprises the phylotypes 8–9 (Table 1).

##### Habitat

The hindgut of higher termites

## 3 Conclusions

To date, the non-methanogenic archaea in termite guts and their potential role in symbiotic digestion have received little attention. Our study provides strong evidence that termite gut *Bathyarchaeia* and other members of the Bathy-6 subgroup are archaeal acetogens: they possess the genomic potential to conserve energy by the production of acetyl-CoA from CO_2_ (*Ca*. Termitimicrobium; TB2) and/or possibly methyl groups (almost all members of Bathy-6, including *Ca*. Termiticorpusculum; TB1). As in bacterial acetogens, their energy metabolism is likely mixotrophic. We identified a complete gene set encoding a novel Fpo-like 11-subunit hydrogenase, which closes the evolutionary gap between the ancestral [NiFe] hydrogenases and the respiratory complex I and would enable members of TB2 to grow lithotrophically on H_2._ All members of Bathy-6 probably derive reducing equivalents from the oxidation of organic substrates (*viz*., amino acids) and might use reductive acetogenesis as an electron sink.

These findings agree with previous claims concerning the capacity for reductive acetogenesis in other subgroups of *Bathyarchaeia*. However, this is the first time that all genes encoding the Wood**–** Ljungdahl pathway and the components required for the provision of reducing equivalents and energy conservation are conclusively documented. Although eight of the nine closely related phylotypes of termite gut *Bathyarchaeia* were represented by high-quality MAGs, a complete pathway was detected only in members of TB2 and two more basal lineages from other environments. This underscores the long-standing caution that the mere presence of marker genes of the Wood**–**Ljungdahl pathway do not qualify an organism as an acetogen since many of its enzymes are are found also in non-acetogenic organisms, where they are involved in the assimilation and interconversion of C_1_ metabolites (Drake, 1994).

## 4 Experimental procedures

### 4.1 Metagenome-assembled genomes (MAGs)

Data on the MAGs from termite guts are from Hervé *et al*. (2020). All other MAGs were retrieved from the NCBI Assembly database (https://www.ncbi.nlm.nih.gov); accession numbers are listed in Table 1. Assembly coverage was determined as described by Hervé *et al*. (2020). Average nucleotide acid identities (ANI) were calculated with fastANI (Jain *et al*., 2018). Protein-coding genes were predicted with Prodigal v2.6.3 (Hyatt *et al*., 2010).

### 4.2 Genome phylogeny

A concatenated gene tree of bathyarchaeotal MAGs was constructed using the deduced amino acid sequences of 43 marker genes extracted with CheckM v1.0.8 (Parks *et al*., 2015). The sequences were aligned using MAFFT v7.305b with the FFT-NS-2 method, and the resulting alignment was filtered using trimAL v1.2 with the gappyout method (Capella-Gutiérrez *et al*., 2009, Katoh and Standley, 2013). Tree topology was inferred with IQ-TREE (multicore v1.6.11; Nguyen *et al*., 2015) using the best-fit evolutionary model suggested by ModelFinder under the Bayesian Information Criterion (Kalyaanamoorthy *et al*., 2017); node support was assessed using the Shimodaira– Hasegawa approximate-likelihood-ratio test (SH-aLRT) with 1,000 resamplings (Lemoine *et al*., 2018).

Taxonomic classification was done with the GTDB-tk version 0.3.2 using the Genome Taxonomy Database (GTDB) release 04-RS89 (https://gtdb.ecogenomic.org/; Chaumeil *et al*., 2018).

### 4.3 16S rRNA gene phylogeny

SSU rRNA gene sequences in the MAGs and other bathyarchaeotal bins obtained from the original metagenomes (Hervé *et al*., 2020) were identified using the *ssu_finder* function implemented in CheckM. Sequences were imported into the alignment of rRNA gene sequences in the SILVA SSURef NR database release 132 (https://www.arb-silva.de; Quast et al., 2013) using Arb v6.0.6 (Ludwig *et al*., 2004). After automatic alignment of the imported sequences using the *PT server* and the *Fast Aligner* tool implemented in Arb, the alignment was manually refined using the Arb editor, considering secondary structure information to identify homologous base positions. After removing sites with more than 50% gaps, the alignment consisted of 1,424 sites with unambiguously aligned base positions. Phylogenetic trees were reconstructed by maximum-likelihood analysis with IQ-TREE using the best-fit evolutionary model (GTR+F+R4) suggested by ModelFinder; node support was assessed using SH-aLRT with 1,000 resamplings. Gene fragments (<1,300 bp) were inserted into the core tree using the *parsimony* tool implemented in Arb.

### 4.4 Gene discovery and annotation

For an initial exploration of the genes potentially involved in energy metabolism, bathyarchaeotal MAGs were analyzed using the annotation provided in the IMG/Mer database (https://img.jgi.doe.gov/mer/; Chen *et al*., 2019). Annotation results were verified, and missing functions were identified with Hidden Markov Model (HMM) searches, using HMMER v3.1b2 (Eddy, 2011) with a threshold E-value of 1E–5; the respective models are listed in Table S3. The identity of all genes of interest was confirmed using the NCBI Conserved Domain search (Marchler-Bauer and Bryant, 2004) and BLASTp (Altschul *et al*., 1990). Additionally, Bathy-6-S, and Bathy-6-B were annotated with BlastKOALA (Kanehisa *et al*., 2016). When indicated, closest neighbors were identified by BLAST, aligned using MAFFT v7.305b with the L-INS-i method (Katoh and Standley, 2013). Phylogenetic trees were reconstructed by maximum-likelihood analysis with IQ-TREE (Nguyen et al. 2015) using the best-fit evolutionary model (LG+G+I) suggested by ModelFinder (Kalyaanamoorthy et al., 2017). Node support was assessed using SH-aLRT with 1,000 resamplings (Lemoine et al. 2018).

### 4.5 Analysis of [NiFe] hydrogenases

Putative [NiFe] hydrogenase genes were identified by HMM searches (see above), using the highly resolved models provided by Anantharaman *et al*. (2016). Search results were confirmed with HydDB, a web-based tool for hydrogenase classification and analysis (https://services.birc.au.dk/hyddb/; Søndergaard *et al*., 2016).

The deduced amino acid sequences of the large subunit (LSU) of [NiFe] hydrogenases recovered from the MAGs and their top BLAST hits on the IMG/Mer database were imported into an alignment of NuoD and FpoD homologs (Lang *et al*., 2015), which was completed with representative members of other hydrogenase classes extracted from HydDB. The alignment was manually refined in the Arb editor. Phylogenetic trees were reconstructed by maximum-likelihood analysis with IQ-TREE (Nguyen et al. 2015) using the best-fit evolutionary model (LG+G+I) suggested by ModelFinder (Kalyaanamoorthy et al., 2017). Node support was assessed using SH-aLRT with 1,000 resamplings (Lemoine et al. 2018).

## 5 Author Contributions

HQL and AB designed the study. HQL analysed data and wrote the first draft of the manuscript. VH contributed to the analyses. AB analysed data and revised the manuscript. All authors edited and approved the final version of the manuscript.

## 6 Acknowledgments

This study was funded by the Deutsche Forschungsgemeinschaft (DFG) in the Collaborative Research Center SFB 987 and by the Max Planck Society. HQL was supported by a doctoral fellowship of the International Max Planck Research School for Environmental, Cellular and Molecular Microbiology (IMPRS-Mic), Marburg, Germany.

The authors thank the Joint Genome Institute for their metagenome sequencing service and for providing the IMG/ER platform.

## 8 Data Availability Statement

The MAGs can be accessed at the NCBI GenBank database (Table 1) and through the IMG platform (https://ncbi.nlm.nih.gov/genbank/ and https://img.jgi.doe.gov/ respectively). Genome IDs and accession numbers for the NCBI SRA database are given in Table S1.

## 11 Supporting Information

**Supplementary Figure S1**. Average nucleotide identity (ANI) of the MAGs in subgroup Bathy-6. The termite gut *Bathyarchaeia* were assigned to phylotypes based on ANI > 99%. NA indicates ANI values <75%, which are not returned by the fastANI program.

**Supplementary Figure S2**. Genome-based phylogeny of termite gut *Bathyarchaeia* illustrating the relationship of lineages TB1 and TB2 to other MAGs in the Bathy-6 subgroup. MAGs mentioned in the text are marked in bold. The maximum-likelihood tree was inferred from a concatenated alignment of 43 proteins using the LG+F+I+G4 model and rooted with selected Crenarchaeota and Euryarchaeota as outgroup. The numbers in circles indicate the phylotypes discussed in the text (Table 1). MAGs included in the comparative analysis (Figure 3) are shown in bold. The tree was rooted other archaeal genomes as outgroup. The scale bar indicates 10 amino acid substitutions per site. Node support values (SH-aLRT) are shown in blue. A simplified version of the tree is shown in Figure 1.

**Supplementary Figure S3**. 16S rRNA-based phylogeny of subgroup Bathy-6, indicating the placement of the sequences from termite guts among those obtained from other environments. The maximum-likelihood tree is based on a curated alignment (1,424 positions) of all sequences in the SILVA database and their homologs retrieved from the bathyarchaeal MAGs (in bold) and the low-quality bins obtained from the termite gut metagenomes (Hervé *et al*., 2020). The tree was rooted using members of Bathy-5 as outgroup. The scale bars indicate 0.05 nucleotide substitutions per site. Node support values (SH-aLRT) are shown in blue. Branches marked with dashed lines indicate shorter sequences that were added using the ARB parsimony tool. A simplified version of the tree is shown in Figure 2.

**Supplementary Figure S4**. The methyltransferase-associated corrinoid protein (CoP) of Bathy-6 and its homologs. (**A**) The canonical methyl transferase system of bacteria and archaea. (**B**) Gene neighborhood of the CoP gene of Bathy-6 and selected homologs (for accession numbers, see panel **C**). Colors indicate the presumed functions of the respective gene products (see panel **A**). Unrooted phylogenetic trees of the methyltransferase-associated CoP genes (**C**) and the associated *mtrH* genes (**D**) of Bathy-6 and their closest relatives (deduced amino acid sequences). Genes that appear in panel **D** are shown in bold. Numbers are IMG/Mer gene IDs. The scale bar indicates 1.0 amino acid substitution per site. Node support values (SH-aLRT) are shown in blue.

**Supplementary Figure S5**. Phylogenetic tree of the catalytic subunit of the Hox hydrogenase of Bathy-6 and its homologs among Group 3 [NiFe] hydrogenases. The maximum-likelihood tree is based on deduced amino acid sequences and was rooted [NiFe] hydrogenase sequences of Groups 1 and 2. The scale bar indicates 0.5 nucleotide substitutions per site. Node support values (SH-aLRT) are shown in blue.

**Supplementary Table S1**. Taxonomic assignment and characteristics of the bathyarchaeotal MAGs from termite guts (from Hervé *et al*., 2020).

**Supplementary Table S2**. Annotation details of the genes that encode the metabolic pathways and other functional markers in the 15 bathyarchaeotal MAGs from termite guts, as discussed in the text (see Figures 3 and 4).

