## Supplementary Figure S1 for "Metabolic potential for reductive acetogenesis and a novel energy-converting [NiFe] hydrogenase in *Bathyarchaeia* from termite guts – a genome-centric analysis"

| 1 | | | 2 | 3 | | 4 | 5 | | 6 | 7 | 8 | | 9 | | | | | |
|---|---|---|---|---|---|---|---|---|---|---|---|---|---|---|---|---|---|---|
| Phylotype_1_Co191P1_bin46 | Phylotype_1_Co191P3_bin4 | Phylotype_1_Co191P4_bin18 | Phylotype_2_Emb289P3_bin80 | Phylotype_3_Lab288P3_bin115 | Phylotype_3_Lab288P4_bin25 | Phylotype_4_Th196P4_bin19 | Phylotype_5_Nc150P3_bin14 | Phylotype_5_Nc150P4_bin1 | Phylotype_6_Cu122P1_bin20 | Phylotype_7_Nt197P4_bin22 | Phylotype_8_Emb289P1_bin127 | Phylotype_8_Emb289P3_bin109 | Phylotype_9_Lab288P3_bin169 | Phylotype_9_Lab288P4_bin61 | Bathy-6-S_SZUA-568 | Bathy-6-B_BE326-BA-RLH | Bathy-6-A_AD8-1 | |
| 100.0 | 99.7 | 99.8 | 81.2 | 80.8 | 80.7 | 78.7 | 78.2 | 78.6 | 78.6 | 79.1 | NA | NA | NA | NA | NA | NA | NA | Phylotype_1_Co191P1_bin46 |
| 99.7 | 100.0 | 99.9 | 81.1 | 81.0 | 81.0 | 78.6 | 78.2 | 78.5 | 78.8 | 79.1 | NA | NA | NA | NA | NA | NA | NA | Phylotype_1_Co191P3_bin4 |
| 99.8 | 99.9 | 100.0 | 81.0 | 80.8 | 80.8 | 78.6 | 78.4 | 78.4 | 78.7 | 79.1 | NA | NA | NA | NA | NA | NA | NA | Phylotype_1_Co191P4_bin18 |
| 81.2 | 81.1 | 81.0 | 100.0 | 82.7 | 82.5 | 79.3 | 78.7 | 78.6 | 79.2 | 80.0 | NA | NA | NA | NA | NA | NA | NA | Phylotype_2_Emb289P3_bin80 |
| 80.8 | 81.0 | 80.8 | 82.7 | 100.0 | 99.6 | 79.5 | 78.6 | 78.8 | 79.2 | 79.7 | NA | NA | NA | NA | NA | NA | NA | Phylotype_3_Lab288P3_bin115 |
| 80.7 | 81.0 | 80.8 | 82.5 | 99.6 | 100.0 | 79.4 | 78.6 | 78.9 | 79.0 | 79.9 | NA | NA | NA | NA | NA | NA | NA | Phylotype_3_Lab288P4_bin25 |
| 78.7 | 78.6 | 78.6 | 79.3 | 79.5 | 79.4 | 100.0 | 78.2 | 78.1 | 78.4 | 78.9 | NA | NA | NA | NA | NA | NA | NA | Phylotype_4_Th196P4_bin19 |
| 78.2 | 78.2 | 78.4 | 78.7 | 78.6 | 78.6 | 78.2 | 100.0 | 99.2 | 80.3 | 78.9 | NA | NA | NA | NA | NA | NA | NA | Phylotype_5_Nc150P3_bin14 |
| 78.6 | 78.5 | 78.4 | 78.6 | 78.8 | 78.9 | 78.1 | 99.2 | 100.0 | 80.5 | 78.9 | NA | NA | NA | NA | NA | NA | NA | Phylotype_5_Nc150P4_bin1 |
| 78.6 | 78.8 | 78.7 | 79.2 | 79.2 | 79.0 | 78.4 | 80.3 | 80.5 | 100.0 | 79.3 | NA | NA | NA | NA | NA | NA | NA | Phylotype_6_Cu122P1_bin20 |
| 79.1 | 79.1 | 79.1 | 80.0 | 79.7 | 79.9 | 78.9 | 78.9 | 78.9 | 79.3 | 100.0 | NA | NA | NA | NA | NA | NA | NA | Phylotype_7_Nt197P4_bin22 |
| NA | NA | NA | NA | NA | NA | NA | NA | NA | NA | NA | 100.0 | 100.0 | 81.4 | 81.6 | NA | NA | NA | Phylotype_8_Emb289P1_bin127 |
| NA | NA | NA | NA | NA | NA | NA | NA | NA | NA | NA | 100.0 | 100.0 | 81.4 | 81.6 | NA | NA | NA | Phylotype_8_Emb289P3_bin109 |
| NA | NA | NA | NA | NA | NA | NA | NA | NA | NA | NA | 81.4 | 81.4 | 100.0 | 99.9 | NA | NA | NA | Phylotype_9_Lab288P3_bin169 |
| NA | NA | NA | NA | NA | NA | NA | NA | NA | NA | NA | 81.6 | 81.6 | 99.9 | 100.0 | NA | NA | NA | Phylotype_9_Lab288P4_bin61 |
| NA | NA | NA | NA | NA | NA | NA | NA | NA | NA | NA | NA | NA | NA | NA | 100.0 | NA | NA | Bathy-6-S_SZUA-568 |
| NA | NA | NA | NA | NA | NA | NA | NA | NA | NA | NA | NA | NA | NA | NA | NA | 100.0 | NA | Bathy-6-B_BE326-BA-RLH |
| NA | NA | NA | NA | NA | NA | NA | NA | NA | NA | NA | NA | NA | NA | NA | NA | NA | 100.0 | Bathy-6-A_AD8-1 |

**Supplementary Figure S1.** Average nucleotide identity (ANI) of the MAGs in subgroup Bathy-6. The termite gut *Bathyarchaeia* were assigned to phylotypes based on ANI > 99%. NA indicates ANI values <75%, which are not returned by the fastANI program.
