## Supplementary Figure S2 for "Metabolic potential for reductive acetogenesis and a novel energy-converting [NiFe] hydrogenase in *Bathyarchaeia* from termite guts – a genome-centric analysis"

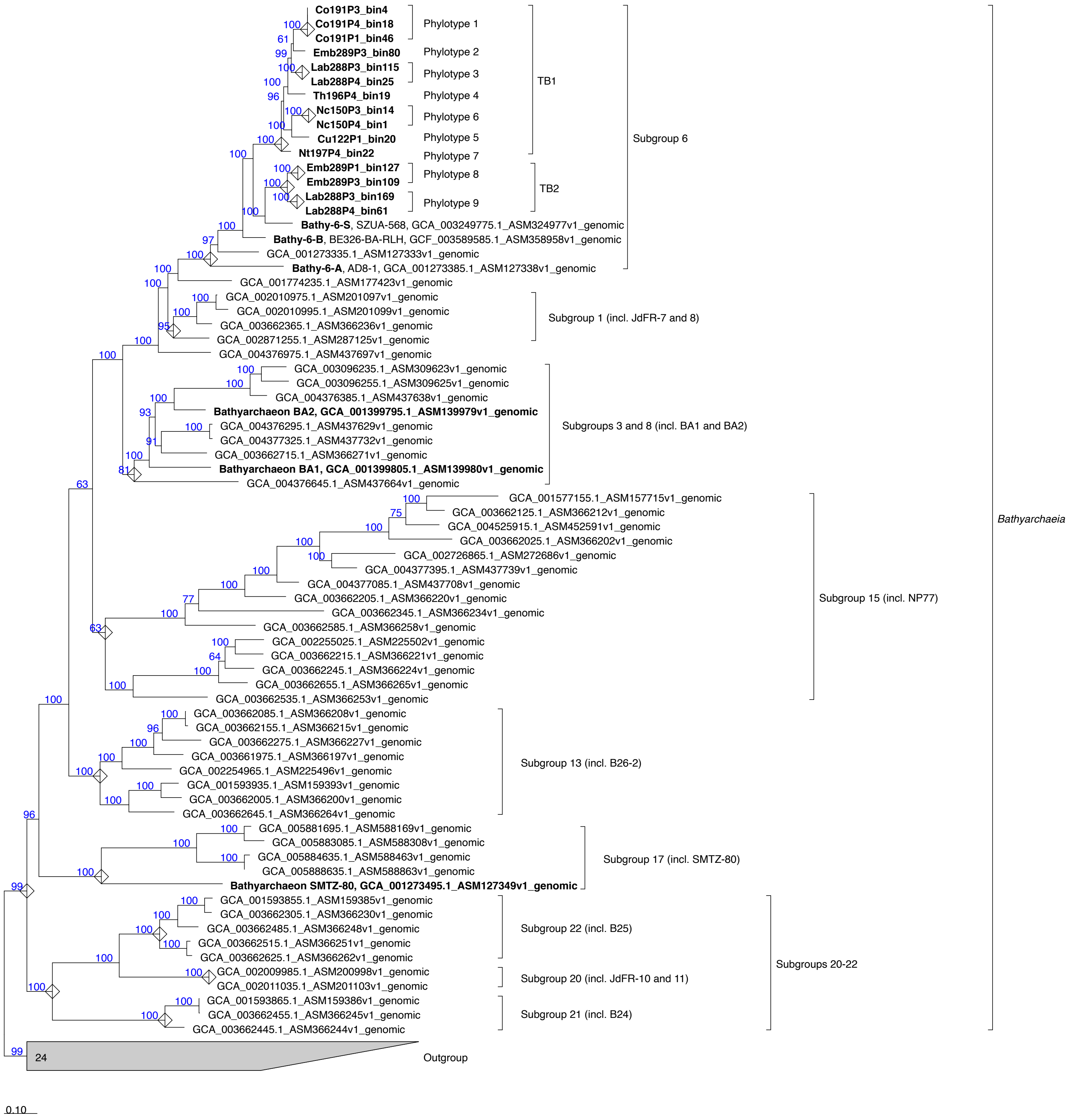


**Supplementary Figure S2.** Genome-based phylogeny of termite gut Bathyarchaeia. The maximum-likelihood tree was inferred from a concatenated alignment of 43 proteins using the LG+F+I+G4 model. The numbers in circles indicate the phylotypes discussed in the text (Table 1). MAGs included in the comparative analysis (Figure 3) are shown in bold. The tree was rooted other archaeal genomes as outgroup. The scale bar indicates 0.1 amino acid substitutions per site. Node support values (SH-aLRT) are displayed at each branch. A simplified version of the tree is shown in Figure 1.
