## Supplementary Figure S3 for "Metabolic potential for reductive acetogenesis and a novel energy-converting [NiFe] hydrogenase in *Bathyarchaeia* from termite guts – a genome-centric analysis"

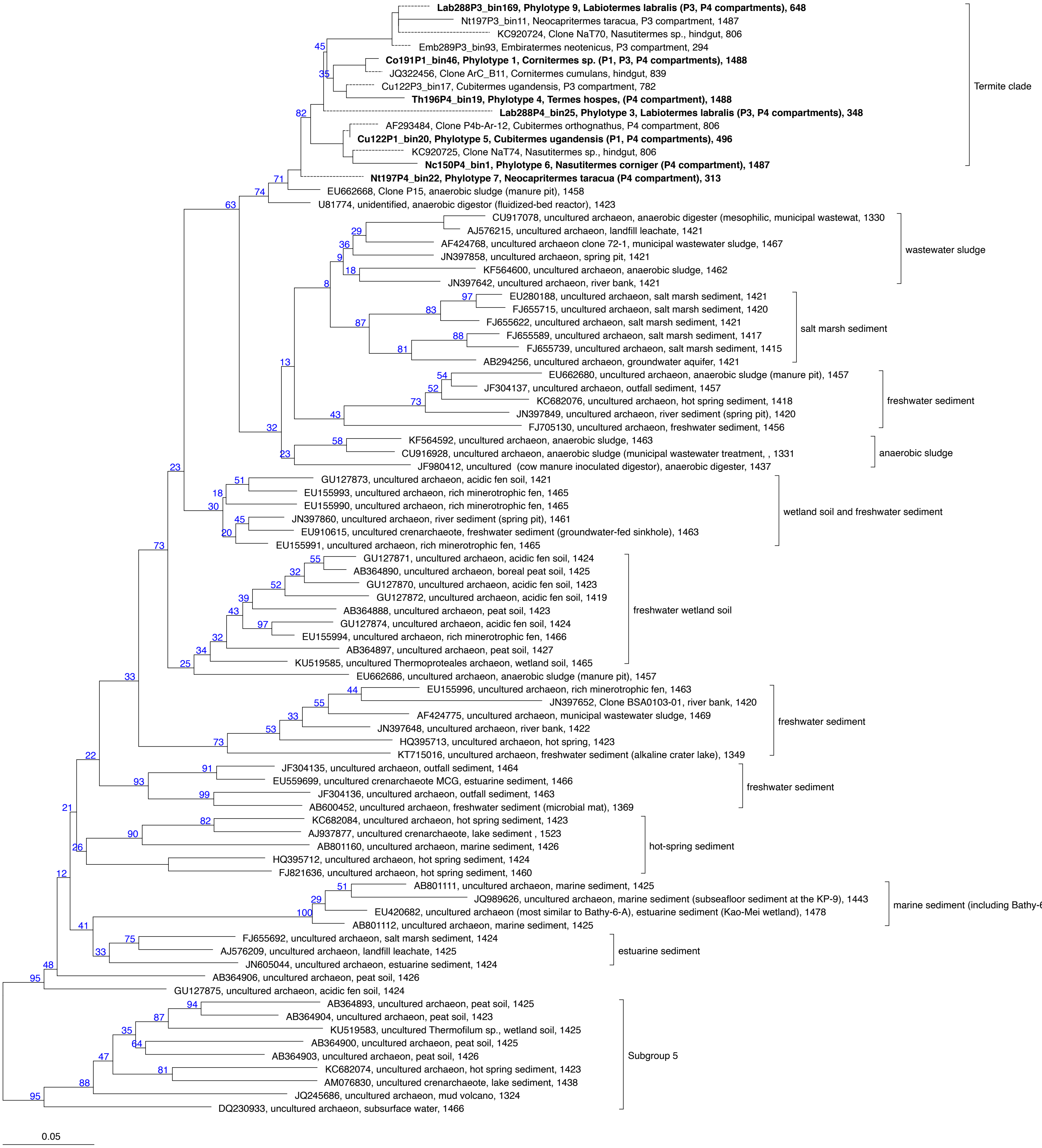


**Supplementary Figure S3.** 16S rRNA gene tree of Bathy-6, indicating the placement of the sequences from termite guts (in bold) among those obtained from other environments. Tree topology was inferred by ML analysis of an unambiguous alignment of the 16S rRNA gene sequences longer than 1300 bp retrieved from the MAGs and also all low-quality bins from the termite gut metagenomes that were classified as Bathyarchaeia (Hervé et al., 2020) in the framework of the SILVA database. The tree was rooted using members of Bathy-5 as outgroup. The scale bars indicate 0.5 nucleotide substitutions per site. Node support values (SH-aLRT) are displayed at each branch. A simplified version of the tree is shown in Figure 2.
