## Supplementary Figure S4 for "Metabolic potential for reductive acetogenesis and a novel energy-converting [NiFe] hydrogenase in *Bathyarchaeia* from termite guts – a genome-centric analysis"

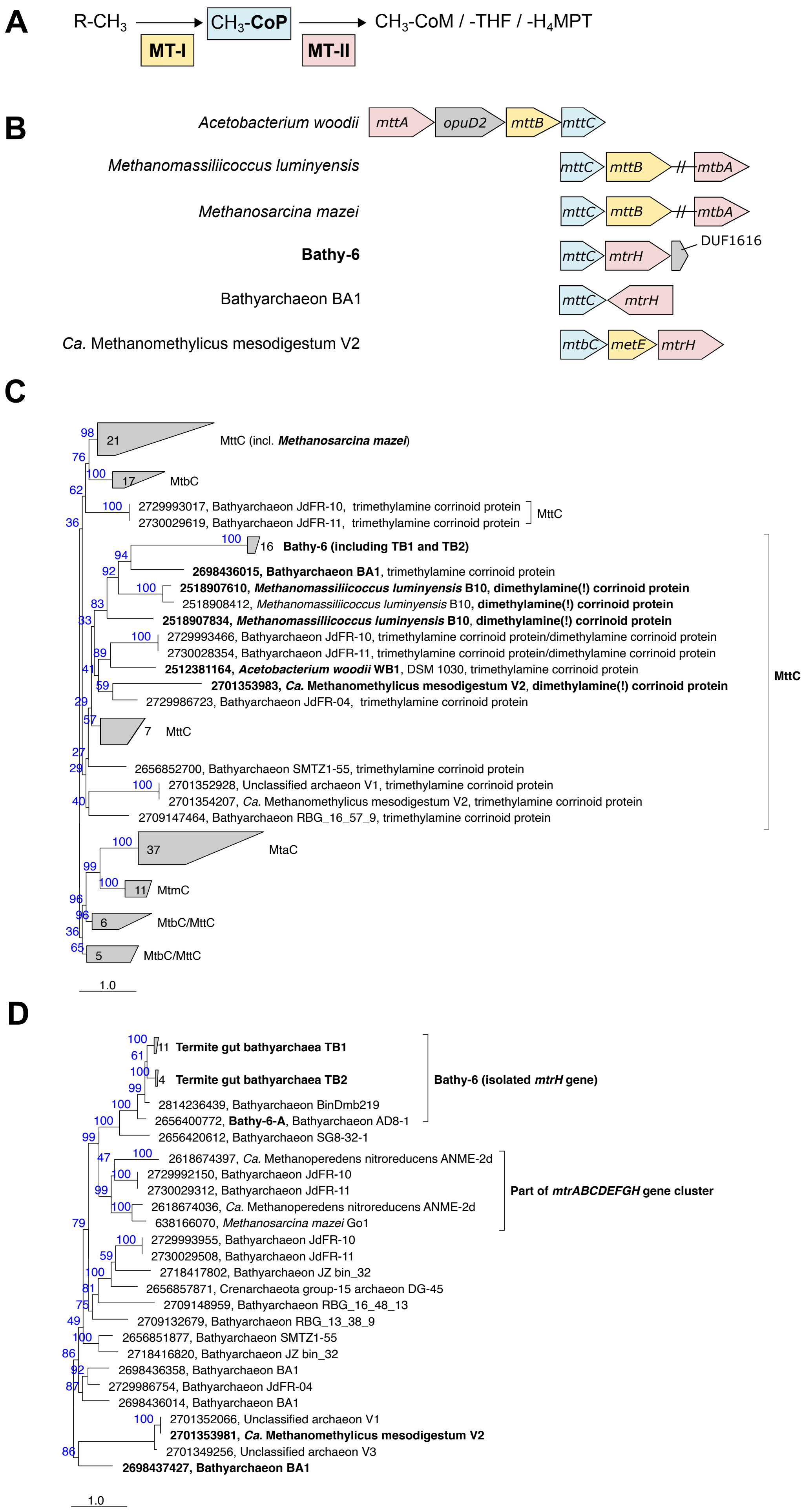


**Supplementary Figure S4.** The methyltransferase-associated corrinoid protein (CoP) of Bathy-6 and its homologs. (A) The canonical methyl transferase system of bacteria and archaea. (B) Gene neighborhood of the CoP gene of Bathy-6 and selected homologs (for accession numbers, see panel C). Colors indicate the presumed functions of the respective gene products (see panel A). Unrooted phylogenetic trees of the methyltransferase-associated CoP genes (C) and the associated mtrH genes (D) of Bathy-6 and their closest relatives (deduced amino acid sequences). Genes that appear in panel D are shown in bold. Numbers are IMG/Mer gene IDs. The scale bar indicates 1.0 amino acid substitution per site. Node support values (SH-aLRT) are shown in blue.
