## Supplementary Figure S5 for "Metabolic potential for reductive acetogenesis and a novel energy-converting [NiFe] hydrogenase in *Bathyarchaeia* from termite guts – a genome-centric analysis"

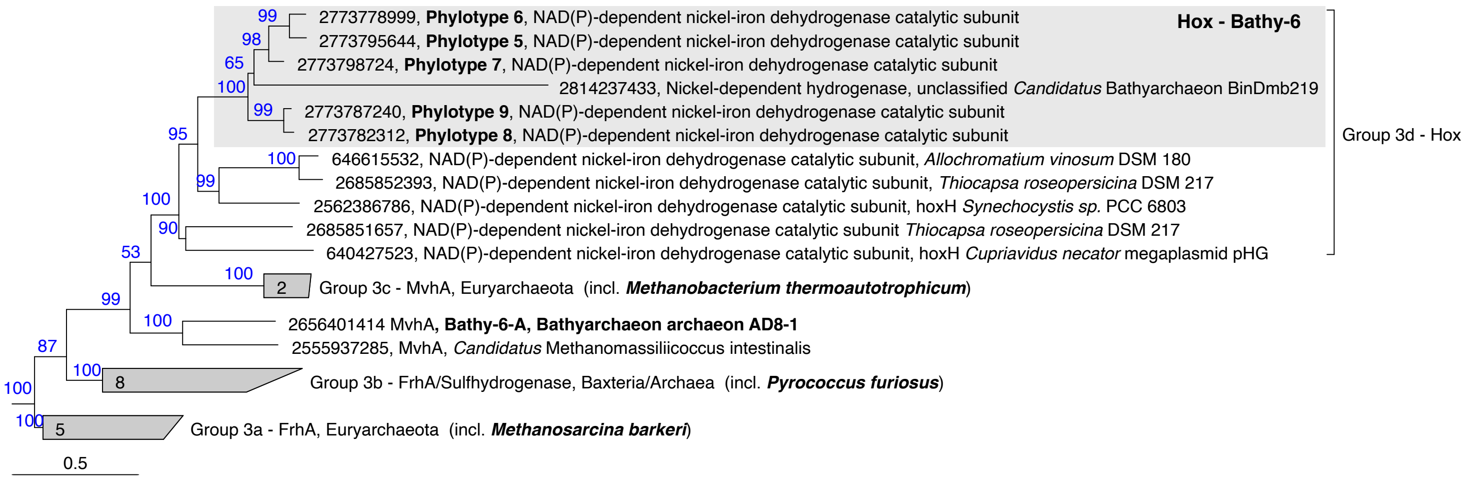


**Supplementary Figure S5.** Phylogeny of the catalytic subunit of the Hox hydrogenase of Bathy-6 and its homologs among Group 3 [NiFe] hydrogenases. The tree was rooted with groups 1 and 2 [NiFe] hydrogenase sequences. Tree topology was inferred by maximum-likelihood; the scale bar indicates 0.5 nucleotide substitutions per site. Node support values (SH-aLRT) are displayed at each branch.
